# Decoding State-Dependent Cortical-Cerebellar Cellular Functional Connectivity in the Mouse Brain

**DOI:** 10.1101/2023.04.11.535633

**Authors:** Yuhao Yan, Timothy H Murphy

**Author notes:** Correspondence to: Timothy H Murphy, Address: 2215 Wesbrook Mall, Vancouver, BC, V6T 1Z3, Canada.

## Abstract

The cerebellum participates in motor tasks, but also a broad spectrum of cognitive functions. However, cerebellar connections with higher areas such as cortex are not direct and the mechanisms by which the cerebellum integrates and processes diverse information streams are not clear. We investigated the functional connectivity between single cerebellar neurons and population activity of the dorsal cortex using mesoscale imaging. Our findings revealed dynamic coupling between individual cerebellar neurons and diverse cortical networks, and such functional association can be influenced by local excitatory and inhibitory connections. While the cortical representations of individual cerebellar neurons displayed marked changes across different brain states, the overall assignments to specific cortical topographic areas at the population level remained stable. Simple spikes and complex spikes of the same Purkinje cells displayed either similar or distinct cortical functional connectivity patterns. Moreover, the spontaneous functional connectivity patterns aligned with cerebellar neurons’ functional responses to external stimuli in a modality-specific manner. Importantly, the tuning properties of subsets of cerebellar neurons differed between anesthesia and awake states, mirrored by state-dependent changes in their long-range functional connectivity patterns. Collectively, our results provide a comprehensive view of the state-dependent cortical-cerebellar functional connectivity landscape and demonstrate that remapping of long-range functional network association could underlie state-dependent change in sensory processing.

## Introduction

Cerebellum has been traditionally thought to mostly contribute to motor coordination and motor learning ^1^. However, empowered by advanced neural circuit interrogation tools, emerging studies in animal models have shown that the cerebellum also plays important roles in higher-order cognitive processes such as spatial navigation ^2^, reward ^3–5^, aggression ^6^ and social behavior ^7^. This is supported by fMRI studies in human subjects where the cerebellum was found to be involved in complex cognitive tasks ^8–10^. Conversely, cerebellar injury or abnormal cerebellum development has been linked to several cognitive and neurodevelopmental disorders such as autism ^11, 12^. This remarkable functional diversity is contrasted by the relative uniform cytoarchitecture and operation principles across the cerebellum ^13^. As such, it has been proposed that the functional diversity of distinct regions of the cerebellum are primarily determined by their differential connectivity to the rest of the brain ^14, 15^.

The cerebral cortex and the cerebellum are two structures that co-evolved through evolution and form one of the largest projection pathways in the brain ^16^. These pathways are characterized by two parts in a closed loop: 1) many areas of the cortex form dense projections to the pons which in turn sends the information to the cerebellum through mossy fibers; 2) Purkinje cells (PCs), the main output neuron of the cerebellum, reciprocally send feedback information to the cortex through the deep cerebellar nuclei (DCN) and the thalamus ^15^. PCs also receive climbing fiber inputs from the inferior olivary nucleus in the brainstem which also receives projections from the cortex ^17^. Due to the indirect nature of these connections, it has been difficult to fully capture the organizing principles of the cortico-cerebellar connectivity, despite its importance in understanding the functional diversity of the cerebellum. Structural connectivity studies using viral tracing techniques provided valuable information on the “hard-wiring” of the cortico-cerebellar connectivity ^18–21^, but may 1) underestimate its complexity due to the fact that these connections are polysynaptic, 2) only provide a post-mortem snapshot of the neuronal connections and miss the dynamic nature of inter-region neuronal networks.

To fill this gap, we correlated mouse cortical population activity recorded using wide-field Ca^2+^ imaging with spiking activity of individual cerebellar neurons, an approach previously shown to effectively map global connectivity patterns of individual neurons ^22–24^. We found cerebellar neurons spanning all recorded lobules that exhibited stable, diverse functional connectivity patterns with the cortex. These long-range connectivity patterns are influenced by local neuronal connections and exhibit state-dependent remapping. Moreover, we demonstrated that cerebellar neurons responding to distinct external stimuli display unique functional connectivity patterns. Notably, we discovered cerebellar neuron ensembles that respond differently to the same stimuli between different states, and this was accompanied by reorganization of their long-range functional connectivity patterns. Our data delineated the spatial-temporal complexity of cortico-cerebellar functional connectivity with cellular resolution and provided a foundation for uncovering the functional identities of individual subcortical neurons by characterizing their global affiliation networks.

## Results

To simultaneously record cerebellar and cortical activity, we employed a Neuropixels probe ^25^, which we inserted at a 45-degree angle from the cerebellum surface, spanning multiple cerebellar lobules. Concurrently, we conducted wide-field calcium imaging of the entire dorsal cortex using an overhead camera in mice expressing GCaMP6 in excitatory neurons (Thy1-GCaMP6s/Emx1-TITL-GCaMP6s^26^). The mice were head-fixed and recorded initially in an anesthetized state for 10 mins using 1% isoflurane, followed by an awake state recording of the same length (Fig.1B). In total, we recorded data from 35 mice, performing 1-4 cerebellar insertions per mouse. The location of each insertion was confirmed through post hoc histology (Fig.1C-D). Fig.1E illustrates the multiunit activity (MUA) of cerebellar units spanning multiple cerebellar layers along the Neuropixels probe during a representative recording.

**Fig.1.**
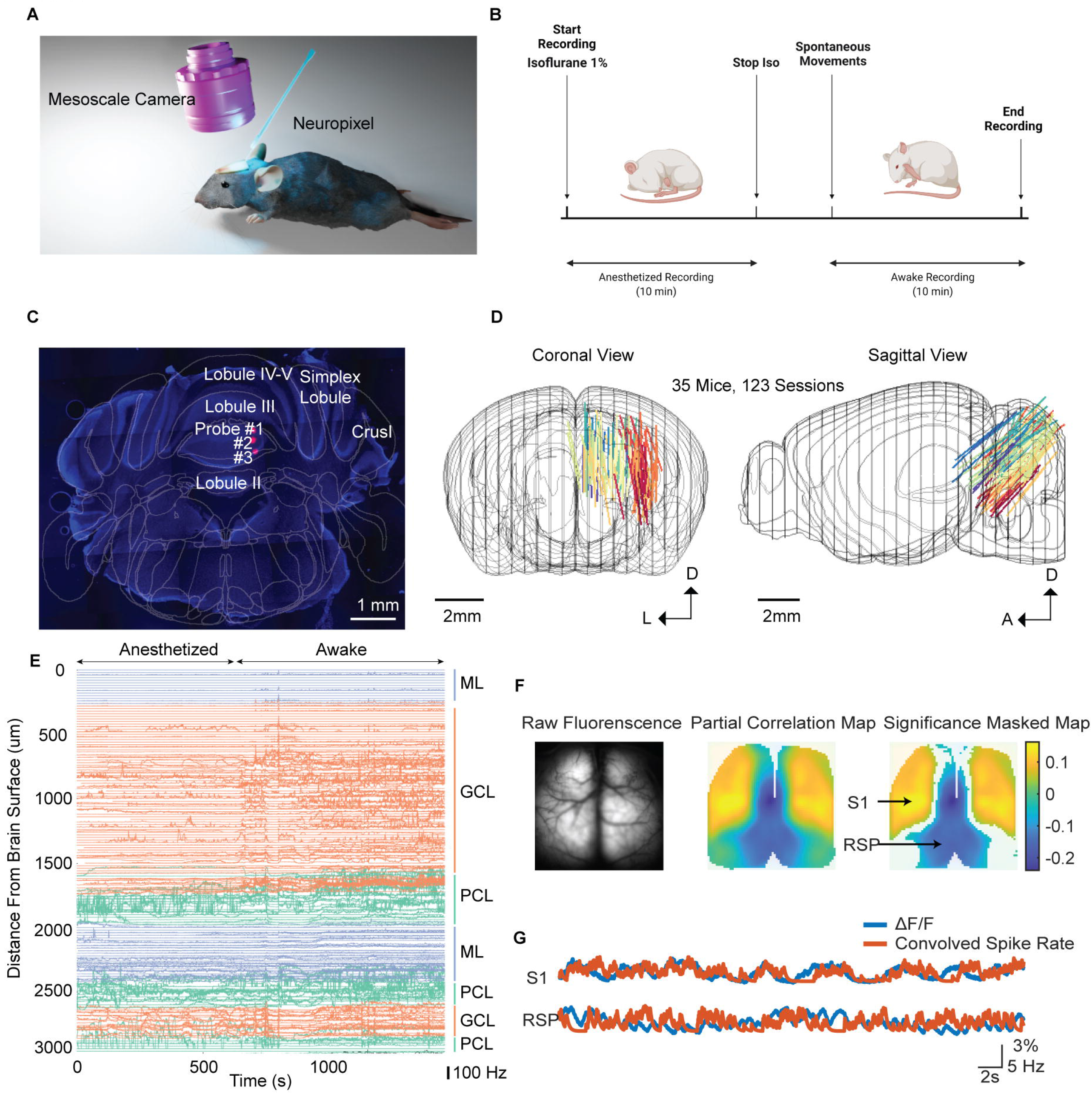
Functional connectivity mapping using in vivo extracellular recording and wide-field Ca2+ imaging. A. Schematic showing experimental setup, with mesoscale epi-fluorescence camera on top of mouse cortex and neuropixel probe in the cerebellum. **B.** Illustration of experiment flow. **C.** Example coronal slice with probe track overlaid with AllenCCF (spatial scale bar: 1mm). **D.** Probe trajectory 3D reconstruction. Each colored dot represents the probe position on a given coronal plane. **E.** Example MUA of cerebellar neurons along the Neuropixel probe color coded by their respective layers. Firing rates were plotted on the same scale as distance, but in Hz. **F.** Functional connectivity maps (CorrMap) of individual cerebellar neurons generated by correlating spiking activity with mesoscale cortical activity with respect to average cortical activity. **G.** Convolved spike train of an example cerebellar neuron overlaid with concurrent ΔF/F traces of the pixel labeled in F.

To characterize the cortical activity network associated with a cerebellar neuron in different behavioral states, we computed two partial correlation maps (CorrMaps) for each cerebellar neuron ^22^, one recorded in an anesthetized state and one in an awake state. Specifically, we calculated the partial correlation value between each pixel time series (ΔF/F) and spike train binned to image acquisition rate (40Hz), with respect to the average activity of the whole cortex to remove the effect of global activity trends. The CorrMaps were used as a proxy for the degree of functional association between a cerebellar neuron and specific cortical regions. To ascertain whether each pixel exhibited a significantly positive or negative correlation with the single-unit activity, we generated a null distribution of correlation values using randomly shuffled spike trains and pixels with correlation value over 95% (positive correlation) or below 5% threshold (negative correlation) were considered significant (Fig.S1D,E, Methods).

To ensure the connectivity patterns shown by CorrMap were stable and not due to spurious activity, we computed the similarity between CorrMaps generated using interleaved epochs and compared that with a null distribution of the epoch similarity generated using shuffled spike trains (Fig.S1B,C, Methods). Only units with these reproducibility metrics that were above the statistically determined thresholds in both anesthetized and awake conditions were included for further analysis. Out of a total of 6699 single units recorded, 2303 units exhibited stable connectivity patterns with the cortex in both anesthetized and awake conditions. Due to the light sensitivity of the Neuropixel probes (and the artifacts induced by strobing), we only assessed the effect of hemodynamic artifacts on CorrMaps in a subset of data (see methods), and found that they have minimal effects on the overall connectivity patterns, consistent with other studies assessing functional connectivity using mesoscale Ca^2+^ imaging ^22, 24^ (Fig.S2). CorrMaps often showed both positively correlated (activated) and negatively correlated (deactivated) regions with opposing functionalities, a hallmark of resting state functional connectivity (rsFC) ^27^ (Fig.1F, G). Additionally, we found that most CorrMaps exhibited high bilateral similarity (Fig.S3), consistent with rsFC patterns observed in mice ^28–30^.

To assess whether the cortical connectivity patterns of cerebellar neurons are more embedded with the cortical network activities or more specific to cerebellar activities, we calculated a library of seed correlation maps (SPM) for each recording and correlated the CorrMap of each cerebellar neuron with the library of SPMs (single seed-pixel maps for all possible pixels) to find the best cortical network that matches each cerebellar neuron’s connectivity pattern. The better the match is, the more the cerebellar neuron’s activity is synchronized with the cortical motif related to a particular seed. SPMs were thought to reflect cortical consensus networks and the underlying intra-cortical synaptic connections, and it has been shown that subcortical neurons often display more specific connectivity patterns that differ from SPMs ^24^. We found there was great diversity to how well CorrMaps of individual cerebellar neurons were matched by the SPMs. For example, in cell#1 shown in Fig.S4, the CorrMap was almost perfectly matched by SPM generated by placing the seed in the left motor cortex, indicating that the spiking activity of this particular cerebellar neuron was highly embedded with the motor cortical motifs. On the other hand, in cell#6, a dominant co-activation of lateral motor cortex, orofacial sensory cortex and visual cortex observed in the CorrMap was not captured by any SPM, highlighting a unique cortical network that was affiliated with this cerebellar neuron (Fig.S4A). Overall, the highest similarities between individual CorrMaps and the corresponding libraries of SPMs were significantly (moderately) increased in the awake state compared to the anesthetized state (Fig.S4C).

## Diverse cortical connectivity profiles of cerebellar neurons

CorrMaps of individual cerebellar neurons exhibited diverse profiles of cortical connectivity patterns, ranging from anterior biased maps which mainly involve the motor and somatosensory areas to posterior cortical maps that involve retrosplenial and visual areas (Fig.2A-C, left to right). To facilitate comparing CorrMaps between different animals, brain images were first parcellated and registered to the Allen Common Coordinate Framework (CCF) using MesoNet, a machine-learning based brain segmentation toolbox ^31^. To capture the diversity of cerebellar CorrMaps, we reduced the dimensional complexity of the CorrMaps by calculating the activation and deactivation indexes for each cell, which is the fraction of pixels that showed significantly positive correlation or significantly negative correlation (Fig. 2B), respectively, with the cerebellar neuron’s spiking activity in each parcellated cortical region ^32^ (Methods). This method has 3 main advantages: 1) compared to using CorrMaps directly as input, activation/deactivation indexes were pre-aligned based on parcellated cortical region contours and easier to compare between animals and sessions, 2) takes into account the number of pixels that were not significantly correlated with cerebellar spiking activity, 3) considers activation pattern of each defined region as a whole and reduces impact of small patches of activation.

**Fig.2.**
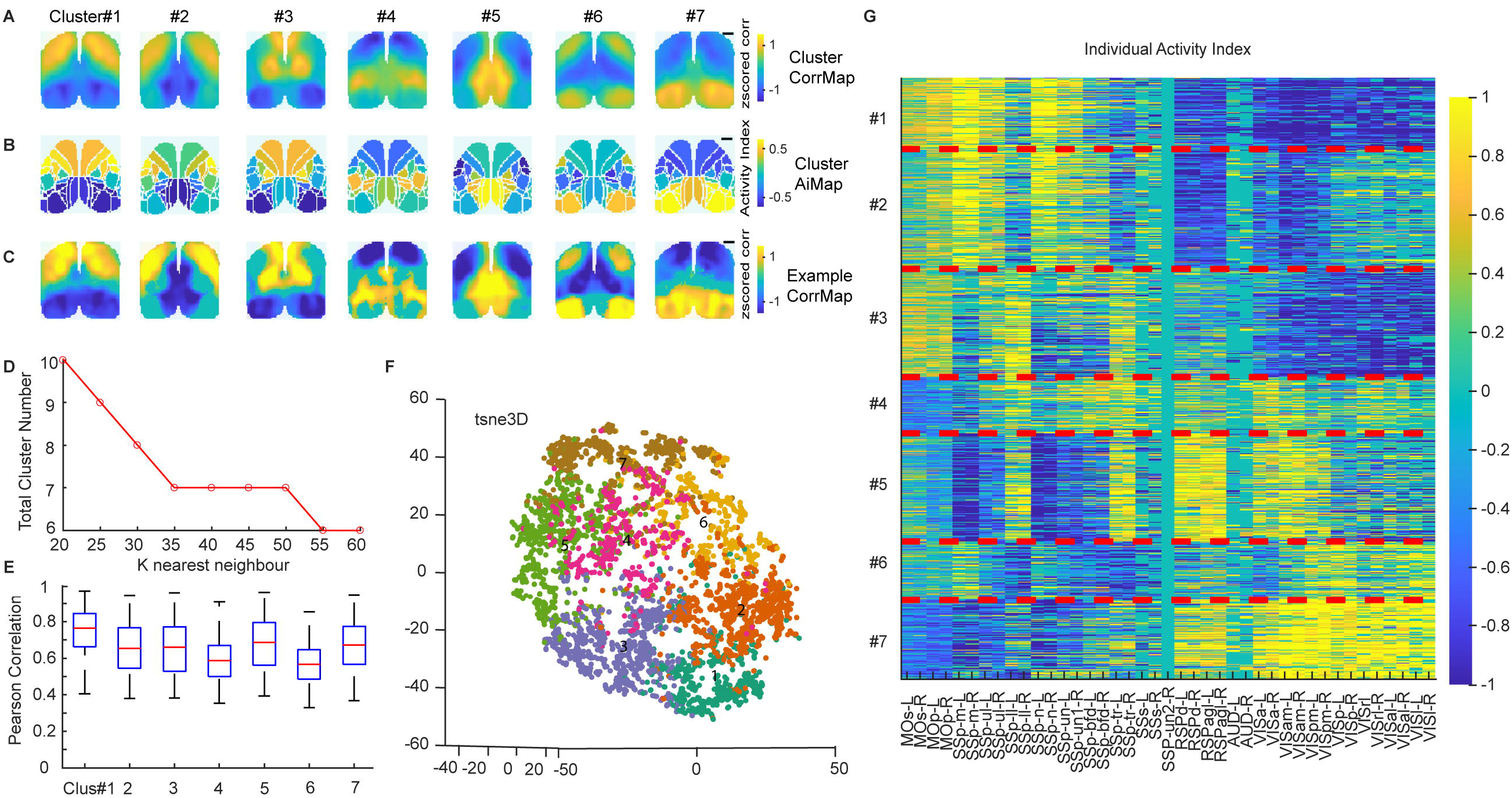
Cluster representation of functional connectivity maps (CorrMap) of cerebellar neurons. A. Average z scored CorrMap of each cluster. Each cluster represents a unique pattern of cerebellar-cortical connectivity network. Spatial scale bar-1mm. **B.** Average activity index map (AiMap) of each cluster. Activity index of each cortical region was computed by summing the activation index vector and deactivation index vector. **C.** Representative individual CorrMap of each cluster. **D.** Resulting number of clusters with varying *k* nearest neighbors used in PhenoGraph clustering algorithm. **E.** Box plot showing correlation between individual CorrMap and average cluster CorrMap. **F.** 3D-tsne plot showing dimensionality reduced representation of clustered CorrMaps. **G.** Clustered activity index of each neuron in the sequence of their cluster assignment. Parcellated cortical regions labeled on the x-axis were ordered in anterior-posterior sequence.

To quantify regional patterns CorrMaps recorded in both anesthetized and awake states from all mice we applied a graph-based clustering algorithm Phenograph ^33^. Clustering led to partition of all cerebellar connectivity patterns into a total of 7 major groups (Fig. 2D, E). In order to validate the optimal number of clusters for partitioning our dataset, we varied k, the number of nearest neighbors used in constructing the weighted graph (see Methods), and found that the standard range of *k* values produced a stable result of 7 clusters (Fig. 2D). Degrees of separation between individual CorrMaps within clusters were quantified (Fig.2E) and visually inspected by projecting dimensionally reduced activation and inactivation indexes onto 3D space using t-SNE (Fig.2F). We then inspected the population connectivity pattern that each cluster represents (Fig.2A-C, G). Fig.2A shows the mean cluster CorrMap, which is the mean of all individual CorrMaps in a cluster. The maps are presented in an anterior to a posterior activated manner, with the individual CorrMap most similar to the cluster average CorrMap shown in Fig. 2C. We also constructed cluster activity index maps (AiMap) by projecting the activity index (sum of activation and deactivation index) of each region onto the reference map (Fig. 2B). Each cluster shows a unique combination of activated and deactivated regions. For example, cluster#1-2 collectively show an activation pattern in the motor cortex while exhibiting varying degrees of activation in somatosensory and visual cortex. Cluster#3 and #4 showed strong activation in the limb and body somatosensory cortex but with motor cortex co-activated in cluster#3 and deactivated in cluster#4. Cerebellar neurons in cluster#5-7, on the other hand, were prominently more correlated with posterior cortical regions, which were thought to be less associated with cerebellar function previously ^21, 34^. Specifically, neurons in cluster#5 had CorrMaps with strong activation in the retrosplenial cortex while the visual cortex was more activated in CorrMaps of cluster#6-7.

## Topographic organization and state-dependent changes in cortico-cerebellar functional connectivity patterns

To assess whether the cortical functional connectivity patterns of cerebellar neurons are topographically organized, we plotted the distribution of CorrMap cluster assignments of neurons in different cerebellar locations across mice. In total we recorded from 8 cerebellar cortical lobules, namely vermal lobule II, III, IV-V, VI, VII, lateral lobule simplex (Sim), CrusI, CrusII, as well as 2 deep cerebellar nuclei (DCN), which are the fastigial nucleus (FN) and dentate nucleus (DN) (Fig.3A-C). Data points from individual animals were only included if they contained at least 10 neurons with stable CorrMaps in both states in a specific cerebellar region.

**Fig.3.**
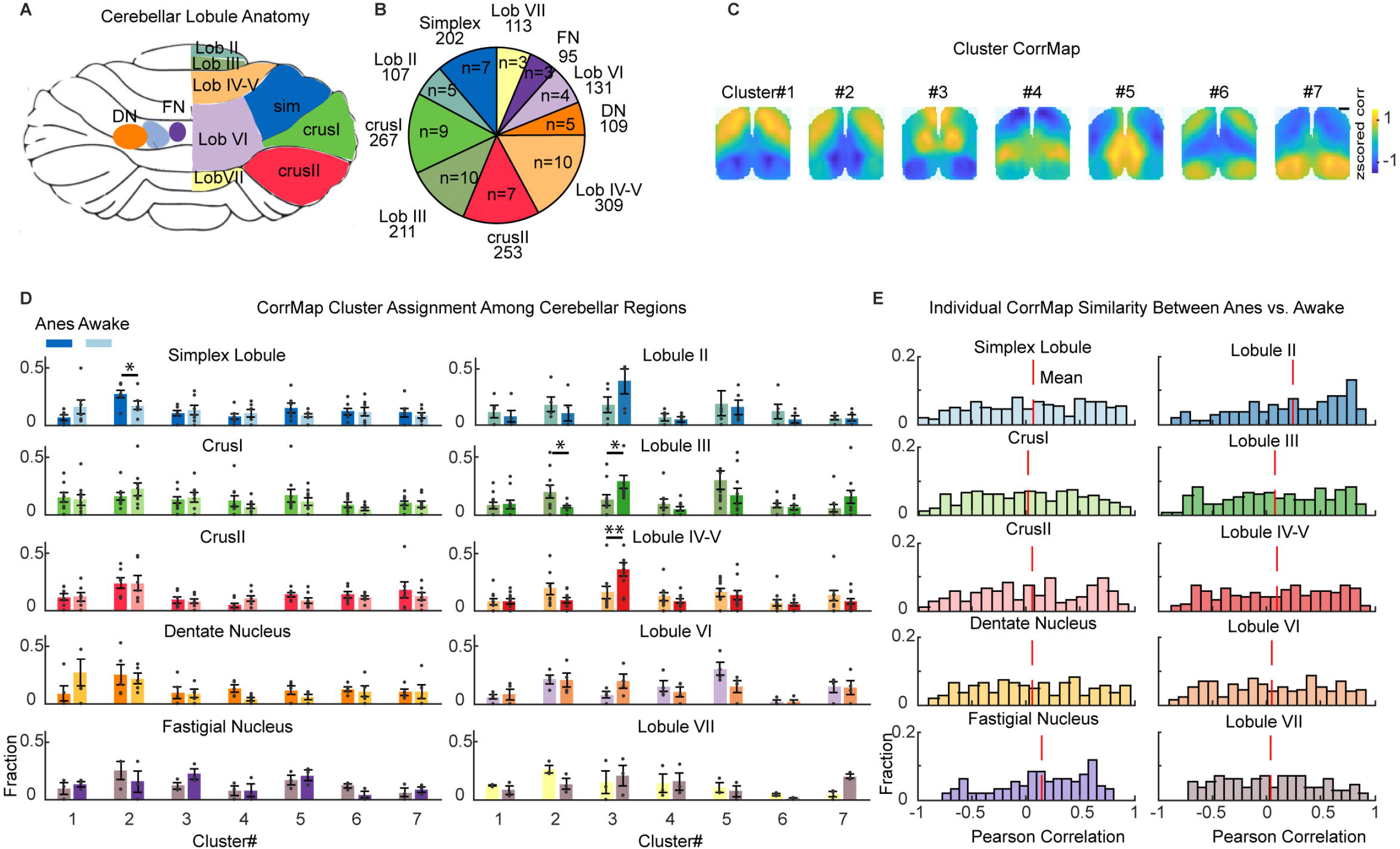
Cerebellar neurons of all regions display diverse profiles of functional connectivity with the cortex during both anesthetized and awake states. A. Schematic showing cerebellar lobules that were recorded from. **B.** Pie charts showing the number of neurons (top number) and the number of mice (bottom number) recorded in each lobule. **C.** Average cortical CorrMap of each cluster for reference. Spatial scale bar-1mm. **D.** The proportion of neurons in each recorded region assigned to each CorrMap cluster during anesthetized (left bar) and awake states (right bar) (Wilcoxon signed rank test, *, **, *** represent p<0.05, p<0.01, p<0.001 respectively). Gray dots represent data from individual mice. **E.** Histogram showing similarity (Pearson correlation value) between CorrMaps of same individual neurons obtained during anesthetized and during awake state. Red dash line shows the mean value of the distribution.

In the anesthetized state, all cerebellar regions (based on neuropixels trajectories) contained neurons exhibiting strong functional connectivity with somatosensory cortex and motor cortex, consistent with structural connectivity profiles of cerebellar neurons reported previously ^18, 21^ (cluster#1-4, Fig.3D left bar). Notably, a number of cerebellar regions also showed substantial functional connectivity (correlational structure) with posterior cortical regions. For example, the lateral cerebellar lobules, namely Sim, CrusI and CrusII all had more than 10% of their neurons that were dominantly affiliated with the posterior visual network (cluster#7, 11.1±3.9%, n=7 mice for Sim; 10.0±2.2%, n=9 mice for CrusI; 17.6±6.9% n=7 mice for CrusII) as well as a considerable fraction that were affiliated with both facial sensory regions and primary visual areas (cluster#6, 12.0±2.8%, n=7 mice for Sim; 8.8±2.4%, n=9 mice for CrusI; 13.6±2.5% n=7 mice for CrusII). Moreover, the anterior vermal lobules exhibited extensive functional connectivity with the retrosplenial cortex (cluster#5, 18.9±11.3%, n=5 mice for Lobule II; 30.1±7.7%, n=10 mice for Lobule III; 15.8±3.2% n=10 mice for Lobule IV-V; 30.1±5.7% n=4 mice for Lobule VI) suggesting that they could play a role in spatial navigation ^2^. Similar to cerebellar cortical regions, both DN and FN contained an appreciable fraction of neurons with cortical representations in all groups. Together, these results indicate that cerebellar neurons with different cortical functional connectivity relationships were dispersed throughout the cerebellum and the long-range functional affiliations of individual cerebellar neurons were not limited by anatomical boundaries within the cerebellum itself ^35^.

Long-range functional connectivity changes between anesthetized and awake states were first evaluated on a population level. We observed that neurons in the anterior vermis showed a consistent increase in fraction of CorrMaps assigned to cluster#3, pattern with activation in motor and limb somatosensory areas (Fig.3C), in the awake state. In particular, neurons in Lobules III and IV-V showed an significant increase in cluster#3 proportion by over 2-fold (Fig.3D, right bar; cluster#3, 12.8±4.7% in anesthetized state to 28.8±5.7% in awake state, p = 0.0195, for Lobule III; 16.3±5.2% to 36.1±6.0%, p = 0.004, for Lobule IV-V; Wilcoxon Signed Rank test). This observation aligns with existing literature suggesting that the anterior vermis plays a crucial role in forelimb reaching tasks ^36, 37^. Moreover, it introduces the possibility that neurons in these regions may adapt to increased involvement in forelimb actions during the awake state by enhancing synchronization with forelimb sensory and motor areas.

In other cerebellar regions, we observed that the overall distribution of CorrMaps stayed mostly consistent between the two states (Fig.3D). However, when we specifically looked at the paired correlation of connectivity patterns of individual neurons across states, we found that almost all lobules exhibited significant variability, even for regions with stable connectivity pattern distribution on a population level (Fig.3E). For example, the overall distributions of functional connectivity patterns in CrusI and CrusII showed little change across states, but the paired correlation of anesthetized and awake CorrMaps of the same neurons in these two regions showed an almost even distribution between -1 and 1 (Fig. 3E). To check that the state-dependent change in CorrMaps we observed was not due to map instability, we recorded an additional anesthetized session following our typical routine in 4 mice (Fig.S5A) and found most neurons showed very stable CorrMaps between the two anesthetized sessions but not the anesthetized and awake sessions(Fig.S5B,C). Thus, the dramatic shift in connectivity patterns of individual cerebellar neurons cannot be explained by instability of their functional connectivity over the recording period, but rather reflect the impact of brain state change on cortico-cerebellar functional connectivity. We also checked whether change in connectivity patterns across brain states was associated with change in firing rates in cerebellar neurons, but found no correlation between the two variables (Fig.S5F). Additionally, spontaneous movements during awake recordings had minimal impact on the CorrMaps (Fig.S6). Taken together, these results suggested that brain state transition can have substantial impact on cortico-cerebellar functional connectivity on the level of individual neurons, but the overall distribution of functional connectivity patterns can remain consistent which may help keep the cerebellum’s capability of processing cortical information of different modalities constant on a population level.

## Cell type-specific distribution and remapping of functional connectivity patterns in mossy fibers and Purkinje cells

We then examined cell type specific distribution of functional connectivity patterns. We focused on mossy fibers (MF) and Purkinje cells (PC) which represent the input and output of the cerebellum, respectively ^38^. Putative MFs were identified based on their stereotypical triphasic and narrow waveforms as well as a narrow gap surrounding 0ms lag in their autocorrelograms whereas putative PCs were identified by their regular firing interval (symmetric harmonics in autocorrelograms) and high firing rates ^39^ (Fig.4B-C, See Methods). Under anesthesia, both MFs and PCs showed populations of neurons with CorrMaps belonging to all 7 clusters. MFs contained lower fraction of neurons associated with visual areas compared to PCs (Fig.4E, cluster#6 and #7 fractions, MF 5.8%±2.4%, 6.4%±1.8%, n = 12 mice; PC 11.6%±1.8%, 14.3%±2.4%, n = 24 mice). In the awake state, MF CorrMaps showed a decrease in fraction assigned to cluster#2 (Fig.4E, anesthetized 25.4%±4.7%, awake 7.7%±1.9%, Wilcoxon signed rank test, p = 5x10^-4^) and an increase in fraction assigned to cluster #3 (anesthetized 18.9%±4.7%, awake 29.8%±5.9%, Wilcoxon signed rank test, p = 0.0273) and #7 (anesthetized 6.4%±1.8%, awake 14.4%±2.2%, Wilcoxon signed rank test, p = 3.9x10^-3^) suggesting a decrease in synchronization with the motor cortex but an increase with somatosensory and visual areas. PC CorrMaps showed a moderate but significant change in proportions of cluster#1 (Fig.4E, anesthetized 7.8%±1.1%, awake 14.0%±2.1%, Wilcoxon signed rank test, p = 0.0152) and cluster#5 (anesthetized 18.9%±2.8%, awake 10.5%±1.9%, Wilcoxon signed rank test, p = 0.0263). Consistent with our previous findings, the paired correlation of anesthetized and awake CorrMaps of individual MFs or PCs exhibited wide dispersion (Fig. 4F), indicating substantial remapping of functional connectivity patterns at the single-neuron level between the two states.

**Fig.4 Cerebellar.**
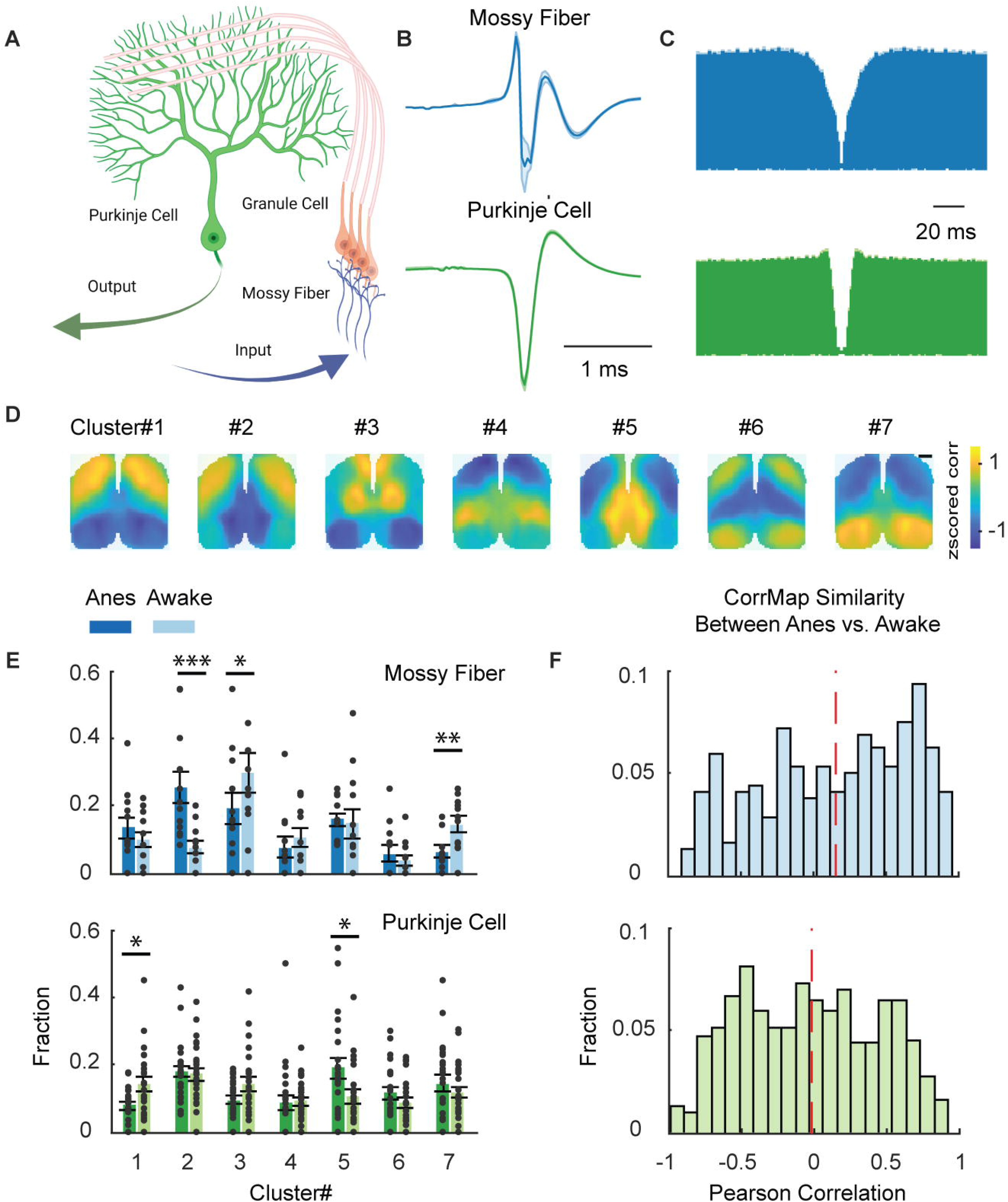
Mossy Fibers and Purkinje Cells exhibited wide distribution of CorrMaps. A. Schematics showing simplified circuitry in the cerebellar cortex. **B.** Averaged waveforms of putative mossy fiber (MF) and Purkinje cell (PC) spikes. **C.** Averaged autocorrelogram of putative MFs and PCs. **D.** Cluster average CorrMap for reference. Spatial scale bar-1mm. **E.** The proportion of CorrMaps of putative MFs and PCs assigned to each CorrMap cluster during anesthetized (left bar) and awake states (right bar) (Wilcoxon signed rank test, *, **, *** represent p<0.05, p<0.01, p<0.001 respectively). Gray dots represent data from individual mice. **F.** Histogram showing similarity (Pearson correlation value) between CorrMaps of same individual MF or PC obtained during anesthetized and during awake state. Red dash line shows the mean value of the distribution.

## Differential encoding of cortical information in simple and complex spikes of cerebellar purkinje cells

Cerebellar Purkinje cells (PC) receive inputs from both parallel fibers (PF) of cerebellar granule cells and climbing fiber (CF) of inferior olive neurons, which give rise to simple spikes (SS) and complex spikes (CS) in PC respectively (Fig.5A). Classic models on cerebellar processing have suggested that there is close interplay between these two input streams where precise conjunctive stimulation of both induces long-term depression (LTD) at the PF-PC synapses, which effectively conveys an error signal that facilitates motor learning ^40, 41^. More recent studies have further highlighted the impact of the context-dependent, information-rich signals carried through climbing fiber inputs on cerebellar circuits and behaviors ^3, 4, 42–44^. However, what specific sensory or behavioral information SS and CS of the same PC represent is less clear. To further delineate the functional roles of SS and CS, we performed pairwise comparisons of CorrMaps of SS and CS of the same PC. We identified SS spike trains of individual PCs and their corresponding CS spike trains based on their cross-correlogram, which would show a hallmark pause of SS firing after CS firing (Fig.5B) if they are indeed from the same PC ^45^. CS with variable waveform shapes were excluded (which suggested that they were recorded from different locations on the PC) to avoid repetitive spike counting. Since CS usually fired at a much lower intrinsic rate, only a small fraction of CS (22/373 for anesthetized CorrMaps, 18/373 for awake CorrMaps) generated statistically meaningful CorrMaps. Interestingly, we observed substantial heterogeneity in similarity of CorrMaps for CS and SS across anesthetized and awake states. Fig.5D and F showed examples where the CorrMaps for CS and SS can either be strikingly similar for some PCs or entirely dissimilar for others. Intriguingly, we observed from individual examples of CS CorrMaps that many showed activation in motor and somatosensory areas. Indeed, when we examined the distribution of all the CS CorrMaps, and we found that the cluster identities of CS CorrMaps were much more skewed towards group patterns with activations in the anterior cortex (clusters 1-3), in contrast to more uniform distribution of SS CorrMaps across all patterns (Fig.5H-I). Taken together, these results suggest that CS preferably encodes sensorimotor information, while SS can encode more diverse modalities of information such that PCs can integrate distinct streams of inputs simultaneously to better facilitate cerebellar function ^46^.

**Fig.5 Cerebellar.**
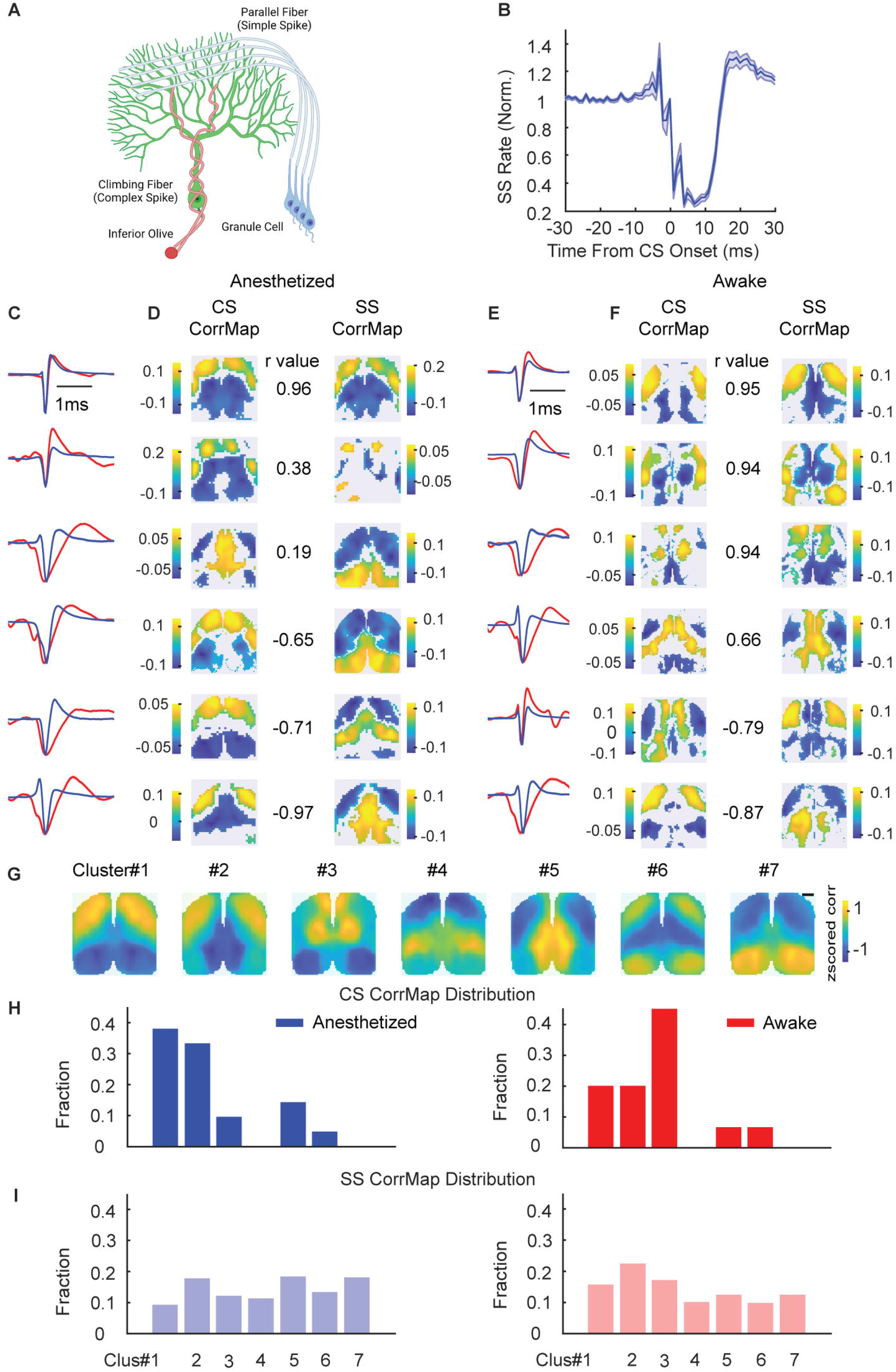
Purkinje cells can encode two distinct streams of functional information from the cortex. A. Cartoon illustration (created using Biorender) of a Purkinje cell receiving inputs from climbing fiber that originates from the inferior olive and from parallel fibers of cerebellar granule cells. **B.** Average cross correlogram of recorded complex spikes and simple spikes of the same Purkinje cell. Simple spikes typically pause for several milliseconds after complex spike firing. **C.** Waveforms of simple spikes (blue) and complex spikes (red) of neurons shown in D. **D.** Purkinje cells displayed a range of similarity between their simple spike connectivity pattern and complex spike connectivity pattern. Numbers between CorrMaps were Pearson correlation values between CS CorrMaps and SS CorrMaps. **E & F.** Same as C & D but showing connectivity patterns in the awake state. **G.** Cluster average CorrMap for reference. Spatial scale bar-1mm. **H-I.** Cluster assignment of CorrMaps of complex spike trains (top) and simple spike trains (bottom) in anesthetized and awake states.

## Local synaptic connectivity influences cerebellar neurons’ participation in distinct global activity networks

Activities of neighboring neurons in highly functionally compartmentalized brain structures such as the cortex and thalamus are often tightly coupled, being potentially shaped by common inputs and recurrent excitation ^22, 24, 47^. Whether this is also apparent for cerebellar neurons, which are less topographically organized by anatomical boundaries in comparison, is unknown ^48^. Since we have found diverse cortical representations correlated with cerebellar neurons residing in all regions we recorded from, we next sought to understand whether the physical distance was a useful predictor of how similar the cortical connectivity patterns of a pair of cerebellar neurons are. To this end, we plotted the physical distances between pairs of cerebellar cortical neurons from the same recording against the similarity of their CorrMaps, and found that there was no relationship between them in both anesthetized and awake conditions (Fig.6A). This indicates that neighboring cerebellar neurons do not necessarily participate in similar distal functional networks, unlike cortical or thalamic neurons where neighboring neurons were usually linked to common activity patterns ^24^.

Considering the heterogeneous organization of cerebellar local circuitries that often transcends anatomical boundaries ^49^, we hypothesized that distant functional connectivity can be more informed by local synaptic coupling than topography. We detected locally coupled cerebellar neuron pairs using cross-correlograms ^50^. Applying a statistically determined threshold, we identified local connections based on peaks or troughs within a +1 ms to 4 ms lag, consistent with monosynaptic interaction timing and the presence of excitatory or inhibitory relationships based on firing probabilities (Fig. 6C, methods). We compared population distribution of CorrMap similarities among putative non-connected (NC), excitatory (EC), and inhibitory (IN) cerebellar neuron pairs with stable CorrMaps; then, we used skewness of the distribution as an indicator of whether CorrMaps were more likely to be similar (left skewness) or dissimilar (right skewness) using un-connected pairs as a reference point. Under anesthesia, NC pairs displayed evenly dispersed CorrMap correlations with slight left skewness, serving as the baseline for CorrMap similarity distribution (Fig. 6F top, Fig. 6G, skewness = -0.177). In contrast, EC pairs showed a significantly left skewness (Fig. 6F middle, Fig. 6G left, skewness = -0.445, Kolmogorov Smirnov test, p = 0.0039, n = 92 and 41,656 pairs for EC and NC, respectively), while IN pairs exhibited a moderate but significant positive skewness (Fig. 6F bottom, Fig. 6G left, blue line, skewness = 0.149, Kolmogorov Smirnov test, p = 0.0257, n = 117 and 41,656 pairs for IN and NC, respectively). This indicated that cortical CorrMaps of EC pairs were more similar to each other, while those of IN pairs were more likely dissimilar (Fig. 6D). In the awake state, the NC pairs’ CorrMap similarities shifted negatively compared to the anesthetized state (Fig. 6F top, Fig. 6G right, skewness = -0.302), resulting in a reduced but still significant left skewness for EC pairs (Fig. 6G right, skewness = -0.471, Kolmogorov Smirnov test, p = 6.2 × 10^-4, n = 188 and 41,656 pairs for EC and NC, respectively). Surprisingly, IN pairs demonstrated a left skewness in the awake state (indicating greater similarity in CorMaps between IN pairs) and were no longer significantly different from the baseline (Fig. 6G right, skewness = -0.286, Kolmogorov Smirnov test, p = 0.3258, n = 187 and 41,656 pairs for IN and NC, respectively). Taken together, these findings suggested that local synaptic connections influence whether cerebellar neurons participate in similar or distinct global activity networks depending on the polarity of the connections (Fig.6B). However, in the awake state, the differences in CorrMap similarity distributions were less pronounced, with the inhibitory pairs no longer significantly different from the baseline (NC pairs).

**Fig 6.**
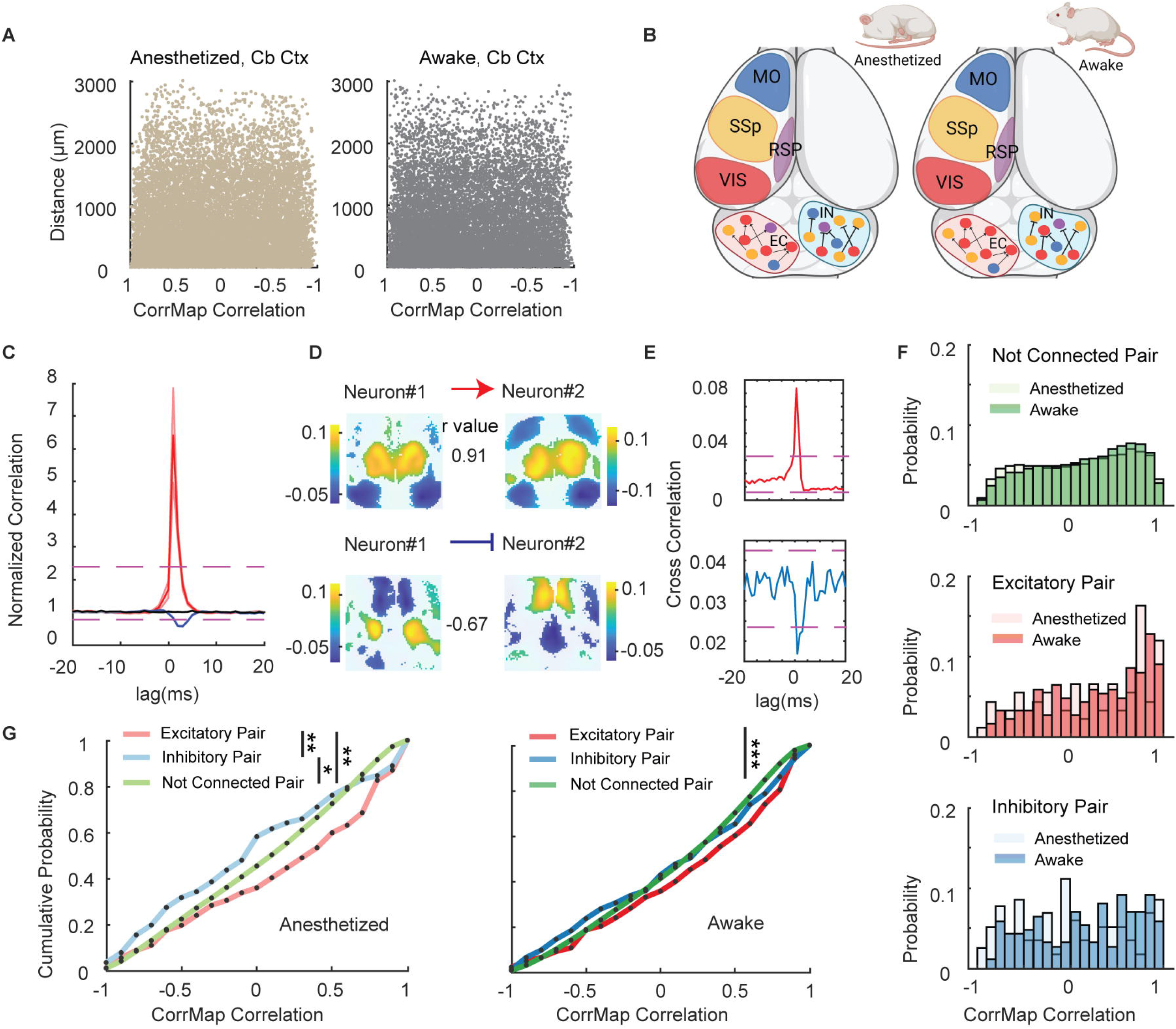
Local synaptic connection but not physical distance between a pair of neurons can indicate the relationship between their long-range connectivity patterns. A. Scatters of similarity between CorrMaps of a pair of neurons in the cerebellar cortex plotted against their physical distance. **B.** Schematic (created using Biorender) showing state-dependent influence of local connections on long-range functional connectivity patterns of cerebellar neurons (with color-coded cortical affiliations). EC pairs were more likely to display similar CorrMaps in both anesthetized and awake states, whereas more IN pairs displayed dissimilar functional connectivity patterns in the anesthetized state but not awake state. **C.** Average cross correlogram of excitatory (red) and inhibitory (blue) pairs of neurons. Null distribution threshold shown in magenta (See Methods). **D.** Example CorrMaps of an excitatory pair and an inhibitory pair of neurons. **E.** Cross correlogram of the pairs in D. **F.** Histogram of similarity between CorrMaps of non-connected (top), excitatory (middle) and inhibitory (bottom) pairs of neurons. **G.** Cumulative distribution function of the CorrMap correlation values between pairs (Kolmogorov Smirnov test, *, **, *** represent p<0.05, p<0.01, p<0.001 respectively).

## Distinct cortical representations of hindlimb and facial stimulation-responsive cerebellar neurons during anesthetized state

Resting-state functional connectivity measured in unconscious states has been suggested to recapitulate relationships within widely distributed task-evoked cortical networks ^51^. Consequently, we investigated whether spontaneous long-range cortical connectivity patterns of individual cerebellar neurons, obtained under our recording paradigm, also contain information on behaviorally-relevant and sensory-related responses. We subjected anesthetized mice to facial and hindlimb stimulation, known sensory stimuli that evoke cerebellar responses ^52–54^, to identify facial stimulation-responsive (FSR) and hindlimb stimulation-responsive (HSR) cerebellar neurons. Following stimulation, we continued our anesthetized recording to compare the spontaneous cortical representations of stimulation-responsive and non-responsive cells.

Stimulation-responsive cells were identified based on normalized z-scored peristimulus time histograms (PSTH) averaged over 40 trials (Fig.7B, see Methods). We only examined neurons that were positively modulated by either stimulus, as a low number of cerebellar neurons were significantly inhibited by facial (24/631) or hindlimb (10/798) stimulation. We confirmed that facial and hindlimb stimulation elicited excitation in the barrel and orofacial sensory cortex, and hindlimb sensory cortex, respectively (Fig.7C, E). Next, we compared the cortical representations of HSR and FSR neurons to non-responsive (NR) neurons by examining the average CorrMap and the mean activity index of each region for the three groups.

**Fig.7.**
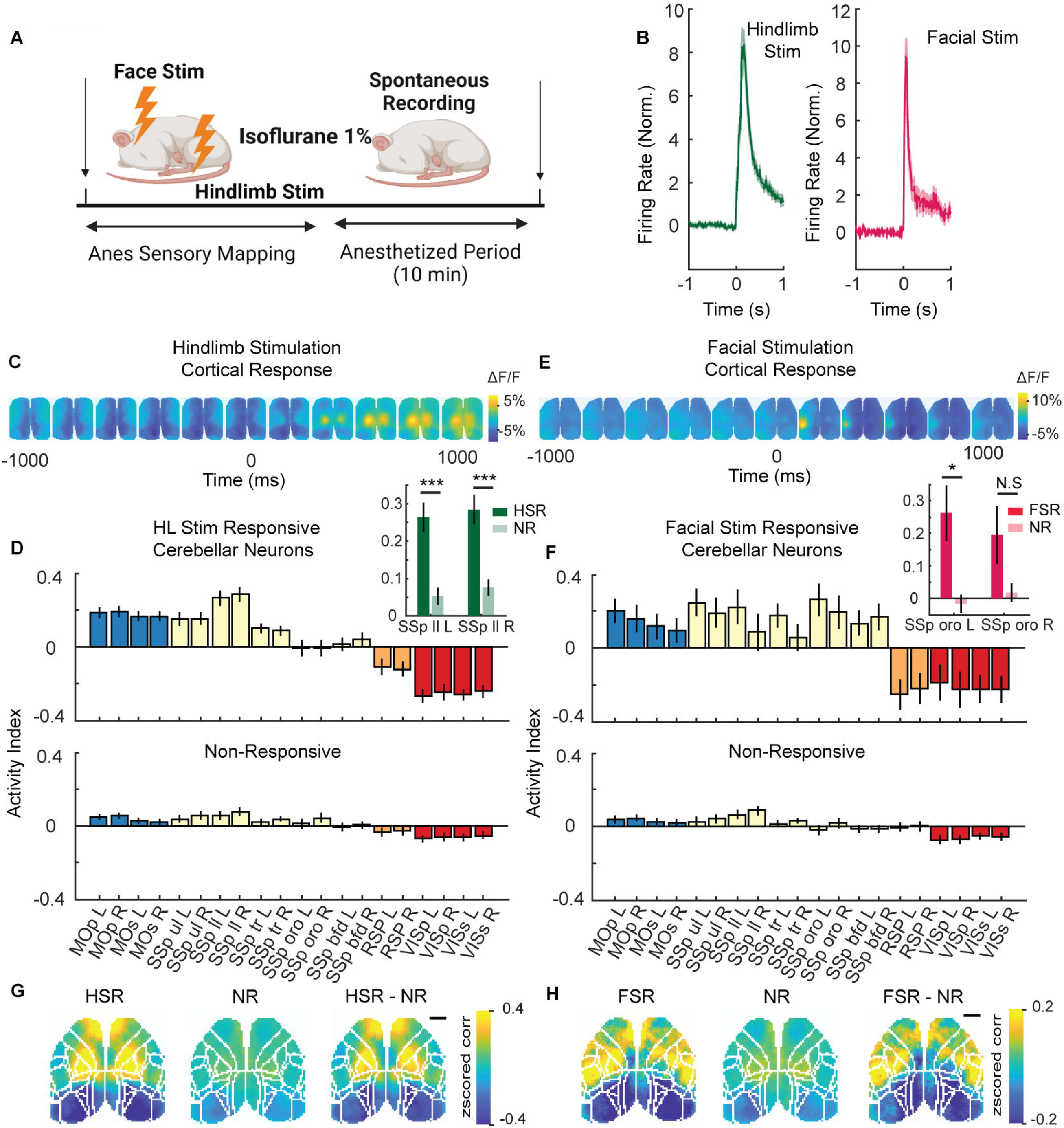
Cerebellar Neurons with Specific Sensory Responses to External Stimuli Displayed Stereotypical Functional Connectivity Patterns. A. Schematic showing experiment flow (Created using Biorender). Under anesthesia, 40 trials of hindlimb stimulation were followed by a 10 mins spontaneous recording from which CorrMaps were calculated. **B.** Average firing rate change in response to hindlimb (HSR) or facial stimulation (FSR) in statistically determined stimuli-responding cells (See Methods). **C & E.** Cortical response to hindlimb and facial stimulation. **D & F.** Average activity indexes of HSR or FSR compared to NR. Insets: Side-by-side comparison of hindlimb sensory cortex or orofacial sensory cortex activity index between HSR or FSR, and NR (Mann-Whitney U-test, *, **, *** represent p<0.05, p<0.01, p<0.001 respectively). **G & H.** Average CorrMaps of HSR or FSR compared to NR, as well as the difference between stimulation response neurons and non-responsive neurons by subtracting their average CorrMaps. Spatial scale bar-1mm.

Notably, HSR neurons’ average CorrMap displayed a distinct pattern compared to NR neurons, with strong activation in the motor and limb somatosensory cortex and deactivation in the posterior cortex (Fig.7G). When we closely examine the cortical region correspond to hindlimb sensory modality, average activity indexes of HSR neurons showed a significant bilateral increase compared to those of NR neurons (HSR, 0.26±0.04, NR, 0.05±0.02, Mann-Whitney U-test, p = 5.93x10^-6^, for SSp ll L; HSR, 0.28±0.04, NR, 0.08±0.02, p = 1.89x10^-6^ for SSp ll R; n = 231 neurons, 15 mice and 639 neurons, 22 mice for HSR, NR, respectively) (Fig.7D, inset). Additionally, FSR neurons’ average CorrMap also differed from that of NR neurons, with activated regions centered around the orofacial sensory cortex (including mouth and nose regions) (Fig.7F). Correspondingly, FSR neurons’ activity indexes revealed a significant increase in the contralateral, but not ipsilateral, orofacial sensory cortex (FSR, 0.26±0.08, NR, -0.02±0.03, Mann-Whitney U-test, p = 0.012, for SSp oro L; FSR, 0.19±0.09, NR, 0.02±0.03, p = 0.159 for SSp oro R; n = 42 neurons, 13 mice and 558 neurons, 17 mice for FSR, NR, respectively) (Fig.7F, inset). These collective findings illustrate that resting-state activity of cerebellar neurons that responded to specific sensory stimuli contained unique cortical representations matching the corresponding sensory modality.

## State-dependent alterations in cerebellar neuron sensory tuning accompanied by changes in functional connectivity with hindlimb sensory cortex

Given this finding, we next sought to investigate whether the state-dependent change in functional connectivity of cerebellar neurons could be a reflection of alterations in their functions, specifically their sensory tuning properties. To this end, we subjected 8 mice to hindlimb stimulation in both anesthetized and awake states followed by spontaneous recording in the two states. We found 23/281 neurons that responded only in anesthetized state but not in awake state (HSR_anes_, Fig.8A, C, D) as well as 64/281 neurons that responded only in awake state (HSR_awake_, Fig.8B, E, F). Interestingly, average CorrMaps of HSR_anes_ neurons indicated a marked decrease in their correlation with hindlimb sensory cortical activity in the awake state compared to the anesthetized state. This was supported by a significant bilateral decrease in the activity index of HSR_anes_ in the hindlimb sensory cortex (Fig.8H inset, anesthetized, 0.36±0.11, awake, 0.07±0.14, Wilcoxon’s signed rank test, p = 0.0424 for SSp ll L; anesthetized, 0.39±0.13, awake, 0.01±0.16, p = 0.0261 for SSp ll R; n = 23 neurons, 8 mice). On the contrary, CorrMaps of HSR_awake_ neurons showed a substantial increase in their functional association with the hindlimb sensory cortex in the awake state compared to the anesthetized state (Fig.8I) paralleled with a significant increase in the activity index of the same cortical region (Fig.8J inset, anesthetized, 0.12±0.08, awake, 0.55±0.07, Wilcoxon’s signed rank test, p = 2.41x10^-5^ for SSp ll L; anesthetized, 0.14±0.08, awake, 0.52±0.08, p = 6.54x10^-4^ for SSp ll R; n = 64 neurons, 8 mice). In summary, spontaneous activity of both HSR_anes_ and HSR_awake_ neurons exhibited enhanced functional connectivity with hindlimb sensory cortex specifically in the behavioral state that they responded to hindlimb stimuli.

**Fig.8.**
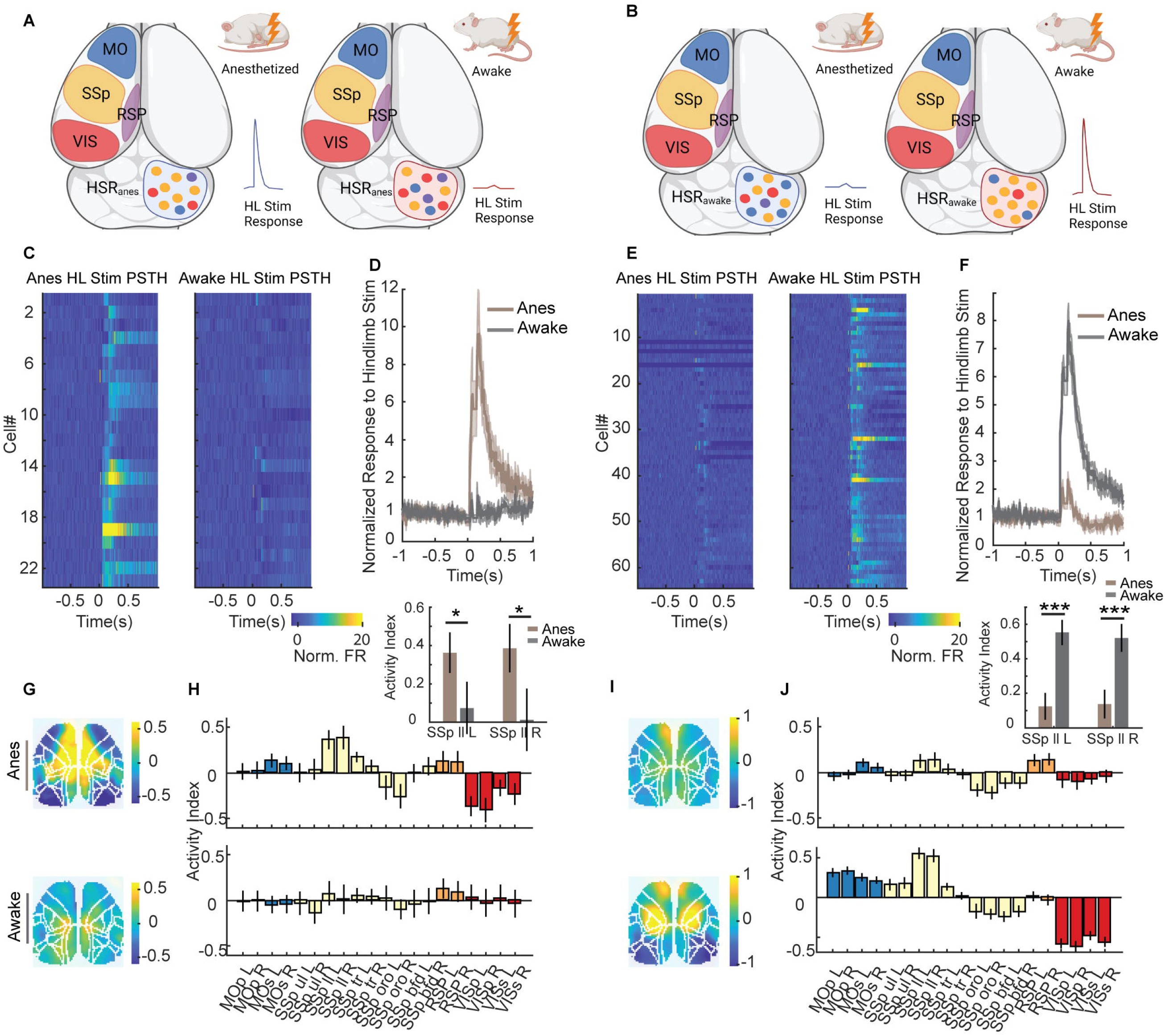
Cerebellar neurons that exhibited change in tuning properties between anesthetized and awake state also showed state-dependent changes in spontaneous functional connectivity patterns. A & B. Schematic (created using Biorender) illustrating separate groups of cerebellar neurons (colors indicate cortical affiliation) that responded to hindlimb stimulation only in an anesthetized (HSR_anes_) or awake state (HSR_awake_) showed strengthened functional connectivity with hindlimb sensory cortex in their respective responsive state. **C & E.** Averaged PSTHs of HSR_anes_ (C) or HSR_awake_ (E). **D & F.** Normalized changes of spike rate in response to hindlimb stimulation of HSR_anes_ (D) or HSR_awake_ (F). **G & I.** Averaged CorrMaps of HSR_anes_ (G) or HSR_awake_ (I) calculated from spontaneous activity in an anesthetized or awake state. **H & J.** Averaged activity indexes of HSR_anes_ (H) or HSR_awake_ (J) calculated from spontaneous activity in an anesthetized or awake state. Insets: Side-by-side comparison between anesthetized and awake hindlimb sensory cortex activity index of HSR_anes_ (H) or HSR_awake_ (J) (Wilcoxon’s signed rank test, *, **, *** represent p<0.05, p<0.01, p<0.001 respectively).

## Discussion

In this study, we combined cerebellar single-unit electrophysiology with wide-field mesoscale cortical Ca^2+^ imaging to refine state-dependent cortical representations that reflect the activity of the cerebellum at cellular resolution. We discovered that cerebellar neurons temporally-linked to diverse cortical representations that were anatomically defined, consistent with intrinsic cortical activity patterns, and dynamically altered between anesthetized and awake states. Cerebellar neurons with similar cortical connectivity patterns did not display topographical organization but rather, local cerebellar synaptic coupling better indicated affiliation with similar long-range functional cortical networks. We also found that both simple and complex spikes of cerebellar Purkinje cells exhibited distinct or linked cortical mesoscale representations, indicating their capacity to integrate multi-modal information. Lastly, spontaneous cerebellar functional connectivity patterns aligned with cerebellar neurons’ functional responses to external sensory stimuli across anesthetized and awake states.

Using high-density recording probes, we captured the heterogeneity of cortico-cerebellar functional connectivity within specific anatomical cerebellar structures. We found a significant fraction of neurons in almost all cerebellar regions are functionally connected to posterior cortical regions, such as PPC, RSP, and V1, beyond traditional associations with motor and sensory cortex ^55^. Previous studies using viral tracing techniques reported only sparse connections between the cerebellum and these posterior cortical regions ^18, 21, 56^. This apparent discrepancy could be because cerebellar neurons’ participation in these networks are multisynaptic and relayed through more nuanced circuitry. Notably, substantial cerebellar projections to the lateral geniculate nucleus (LGN) of the thalamus suggest cerebellum’s functional involvement in visual processing networks, despite weak direct connections to the visual cortex ^21^. Functional studies on mice with impaired plasticity in parallel fiber-Purkinje cell (PF-PC) synapses showed selective impairment in hippocampal place cells, indicating the cerebellum’s role in shaping spatial representations, a key RSP function ^2^. Our results support a previously underrecognized functional network connecting cerebellar regions with the posterior cortex. Further research is needed to elucidate how cerebellar neurons participate in cortical functions, such as decision-making, spatial navigation, and visual processing.

We discovered that diverse cortical representations associated with individual cerebellar neurons can undergo significant remapping when animals transition from anesthetized to awake states. While some cerebellar regions showed relatively unchanged overall state-dependent cortical representations, individual neuronal representations within these areas varied across states (Fig.3D, E). This suggests that long-range communications between cortical and cerebellar regions remain consistent on a population level, but individual neurons’ functional identities are dynamically regulated in a state-dependent manner. Our results align with research indicating that functional connectivity between brain regions is dynamically shaped in state- and behavior-dependent ways to accommodate varying tasks and environments ^22, 32, 57^.

Unlike the topographically organized cortex ^29^, the cerebellum’s fundamental processing modules, or microzones, are sagittally organized functional zones that are better differentiated by genetic markers and climbing fiber input origins of PCs ^46, 58^. PCs in different microzones engage in distinct behavioral tasks, attributed to their specific intrinsic properties and input/output pathways ^48, 59^. Our study presents a unique opportunity to examine how cerebellar neurons’ activity and long-range connectivity patterns are coordinated within the same sagittal plane. Our findings reveal that cerebellar neurons within the same sagittal plane exhibit diverse functional connectivity patterns, and neighboring neurons do not necessarily participate in similar long-range networks, similar to what was observed in cortical neurons ^32^. This suggests more complexity in how cerebellar neurons’ geometry relates to their functions. Moreover, we found that locally coupled cerebellar neurons are more likely to engage in similar or dissimilar networks for excitatory or inhibitory synaptic connections, respectively. This implies that functionally clustered local circuitry forms consensus affiliations with long-range cortical networks, with multiple local networks potentially arising within the same microzone to facilitate complex information processing. Future experiments combining wide-field cortical and cerebellar imaging across multiple sagittal planes will clarify cortico-cerebellar connectivity organization between different cerebellar microzones.

One salient feature of the PCs in the cerebellum has been the unique organization of their afferents, namely thousands of parallel fibers from granule cells and a single climbing fiber from inferior olivary neurons that each PC receives ^41^. The interaction between these two streams of inputs and how they converge to induce synaptic plasticity mechanisms in PCs has been central to our understanding of cerebellar function in motor control and learning. Here we found that complex spikes in most PCs we recorded are functionally affiliated with the anterior cortex, suggesting that they may mainly carry motor or somatosensory signals. On the other hand, the simple spikes of these PCs exhibit a wide variety of cortical representations, ranging from sensorimotor cortex related to retrosplenial or visual cortex related. A plausible model stemming from these observations is that CSs encode specific feedback sensorimotor signals that are integrated with multimodal signals encoded by SS spikes that cooperatively facilitate motor learning. Emerging literature has started to delineate the involvement of complex spikes in higher-order reinforcement learning processes ^3, 4, 60^. A compelling direction for future research would be to investigate the extent to which cortical representations of complex spikes can adapt beyond sensorimotor signals following the acquisition of task rules that associate previously unrelated stimuli ^61^. Further experiments should address the dynamics of feature-encoding by multi-regional complex and simple spikes in the context of cerebellum-specific synaptic plasticity rules under various task learning paradigms.

The biological significance of resting state functional connectivity has been debated ^62, 63^. Previous studies have shown that connectivity patterns from spontaneous activity can predict task-evoked or stimulus-evoked responses in brain regions ^51^ or even single neurons ^64^ but most of these investigations focused on intracortical connectivity. Extending this approach to explore subcortical regions’ unknown physiological functions is a logical step. We demonstrated that analyzing functional connectivity is useful for understanding individual cerebellar neuron functions. Specifically, cerebellar neurons that responded to facial or hindlimb stimulation showed distinct cortical representations under spontaneous and anesthetized states, reflecting unique information about their functional identities. We further showed that cerebellar neurons displayed state-dependent changes in their sensory responses which was mirrored by a concerted change in their functional connectivity patterns between states. This result suggests that alterations in long-range communication across brain structures could be a novel mechanism underlying functional differences between brain states ^65^. This also leaves the question of what is the functional significance of the information flow between cerebellum and higher order cortical areas such as PPC and prefrontal cortex, as these areas were dominantly activated in CorrMaps of some cerebellar neurons. Given the substantial improvement in ease of access and throughput of electrophysiological recordings of subcortical neurons enabled by advances in recording probes ^66^, the combined cortical wide-field imaging and single neuron recording approach can be extended to characterize cortical mapping of other subcortical regions and resolve their functional identities at cellular resolution.

## Methods

### Animals

All animal use and experimental procedures were in accordance with animal protocols (A18-0036) approved by the University of British Columbia Animal Care Committee and Canadian Council on Animal Care. Mice were group-housed in a vivarium on a 12 hr day light cycle (7 AM lights on). To achieve wide-field imaging of cortical activity, transgenic mice expressing GCaMP6s in cortical excitatory neurons were used. These included Thy1-GCaMP6s strain ^67^ (JAX #024275, https://www.jax.org/strain/024275) and Emx1-TITL-GCaMP6s strain produced by crossing Emx1-Cre mice (JAX#005628, https://www.jax.org/strain/005628) with Camk2a-tTAxAi94(TITL-GCaMP6s) mice ^26^ (obtained from Allen Institute). Data were collected from 19 male and 16 female mice ranging between P40 and P80. Sample sizes were not predetermined but consistent with other studies on similar topics ^22, 24^. No clear functional difference was found between mice of both sexes and subjects were pooled and analyzed together.

### Surgery

Mice underwent surgical procedures to prepare for widefield imaging and cerebellar recording on the same day of the recording sessions. During surgery, mice were induced under anesthesia using 1.5%-2% isoflurane which was then reduced (0.5%-1%) for maintenance. The scalp of the animal was removed to expose the skull covering the olfactory bulb, dorsal cortex and the cerebellum, which was then fixed to a custom-made stainless steel headplate using cyanoacrylate glue and dental cement. A 3mmx3mm craniotomy was then made on the right cerebellum to prepare for probe insertion. Throughout the surgery and recordings, body temperature of the animal was monitored and maintained at 37L using a heating pad with a feedback thermistor.

### Electrophysiological Recordings

Spiking data of cerebellar neurons were recorded using Neuropixels probe ^25^(Phase 3B) which was mounted on a custom 3D-printed rod adaptor. The rod adaptor was fixed on a motorized micromanipulator (Sutter instrument, MP-225) parallel to the midline of the brain and at a 45° angle relative to the horizontal plane. The probe was inserted parasagittally into the cerebellum crossing multiple lobules to a depth of 3-3.5mm. All insertions were made on the right cerebellum. The probe was coated with lipophilic dye (DiI, ThermoFisher Cat No. D3911) for post-hoc histological analysis. External reference made from a copper wire soldered to the probe was used. Recordings were acquired at 30kHz with gain of 500x and on-line application of a high-pass filter (300Hz) using OpenEphys ^68^ (https://github.com/open-ephys)

Single units were identified first through automatic spike sorting software Kilosort2.5 ^69^ (https://github.com/MouseLand/Kilosort) and followed by manual curation using Phy2 ^70^ (https://github.com/cortex-lab/phy). Units with contamination percentage>10% (calculated by dividing the event rate within 2ms bin of unit autocorrelogram by the max height of autocorrelogram), mean spike rate<0.3Hz, missing spikes during significant periods of the recordings and amplitude<40µV were excluded from analysis.

Purkinje cells were identified through their typical high firing rate, regular firing pattern and presence of complex spikes ^71^. Simple spikes and complex spikes were attributed to the same Purkinje cell if simple spike firing showed a typical pause following a complex spike by inspecting their cross-correlogram (Fig.5B). Waveforms of complex spikes showed either a large narrow trough followed multiple spikelets or a broad trough, an indication of the recording site being perisomatic or dendritic, respectively ^45^ (Fig.5C, E), and complex spike trains exhibiting both waveforms were excluded to avoid repetitive spike counts.

### Widefield Ca^2+^ imaging

Widefield images of dorsal cortical activity were acquired using a pair of front-to-front video lenses (50 mm, 1.4 f:30 mm, 2 f) coupled to a 1M60 Pantera CCD camera (Dalsa) ^24^. During recording, the surface of the dorsal cortex was evenly illuminated with blue light emitted from a blue-light-emitting diode (Luxeon, 470 nm) coupled with a bandpass filter (467–499 nm). The emission fluorescence from the calcium indicator GCaMP6s was filtered by a 510–550 nm bandpass filter. 12-bit images were collected at a frame rate of 40Hz during widefield recording using XCAP imaging software. These images were binned by a factor of 8 which produced final images with 128x128 pixel dimensions with ∼68µm per pixel. The blue light did have a constant impact on absolute voltage values of Neuropixels probe recording, but the impact was inconsequential as it was removed by common average referencing during analysis.

To address the impact of hemodynamic response on global connectivity patterns of cerebellar neurons, we performed strobed recordings for a subset of the data where 470nm blue light and 530nm green light were turned on in an alternating fashion at 80Hz (to keep the same frame rate for brain activity recordings). 530nm green light is at the isosbestic point of the oxy/deoxy-hemoglobin absorption spectra and thus the green reflectance signal can capture brain blood volume changes ^72^. Strobed recordings were not performed on all datasets since alternating light caused artifacts in Neuropixels recordings. We found hemodynamic correction had minimal effects on the overall connectivity patterns of cerebellar neurons, consistent with results reported previously ^22, 24^.

### Awake Recordings

Simultaneous wide-field imaging and electrophysiological recordings were first performed for 10 mins when the animal was under light anesthesia (1% isoflurane). Then, the isoflurane flow was turned off and the animal was allowed to wake up for at least 3-5 mins. Spontaneous movements made by the animal were considered signs of consciousness and awake recordings were subsequently made for 10 mins. For a subset of mice, videos of the mice were taken during the awake recording. Optical flow of the mouse body (MATLAB estimateFlow function) was used to detect periods where the mouse was making spontaneous movements; timepoints where optical flow was greater than 2 standard deviation plus mean were considered movement periods. Quiet periods referred to periods where the animal was not moving.

### Histology

After recordings, mice were subject to transcardial perfusion with 4% paraformaldehyde (PFA) dissolved in standard phosphate-buffered saline (PBS) solution. The dissected brain was submerged in 4% PFA-PBS solution at 4°C overnight, before being cut into 100µm sagittal or coronal slices using a vibratome (LEICA VT1000S). Slices were subsequently laid flat on glass slides (Fisher Scientific Cat No. 12-550-15) and mounted with DAPI-fluomount (SouthernBiotech Cat No. 0100-20). Histological images were acquired using Zeiss inverted microscope (Zeiss Observer z1) accompanied with Zen Pro software. DiI track was used to confirm probe locations. Probe tracks were reconstructed using open source code (https://github.com/petersaj/AP_histology) and registered to the Allen Mouse Brain Common Coordinate Framework ^73^ (Allen CCF). The recording channel corresponding to brain surface was identified using sudden drop of LFP power between in-tissue and out-of-tissue recording sites (https://github.com/AllenInstitute/ecephys_spike_sorting).

### Sensory Stimulation

For facial and hindlimb stimulation, thin acupuncture needles (0.14mm) were implanted ipsilaterally (right side) into the facial muscle around the nose area or the hindpaw, respectively. 1.5mA, 50ms stimulation current was generated using a stimulation isolator (World Precision Instruments A385) coupled with a pulse stimulator (A-M Systems 2100) and used to stimulate through the needles. Cortical and cerebellar response to sensory stimulation was calculated from averages of 40 trials with 5s inter-stimulus interval. Stimulation pulse was recorded using OpenEphys software and synced to electrophysiological and wide-field recordings.

### Data Analysis

#### Wide-field Imaging Data Preprocessing

Wide-field images were binned by a factor of 2 (end results 64x64 images) by MATLAB nearest-neighbor interpolation method. Δ*F/F* was considered as a proxy of cortical activity and was calculated by first subtracting baseline value from each frame then dividing by the same baseline. Baseline value was calculated as the median fluorescence value of each pixel 20s preceding any given time point. In datasets where hemodynamic response were corrected, Δ*F/F* of the green reflectance images were separately calculated and subtracted from the brain activity Δ*F/F*.

#### CorrMaps Construction and Quality Control Measures

We adopted a previously published method ^22^ to find correlational relationship between cortical activity and spiking activity of individual cerebellar neurons, which we used as the proxy for the cortical functional connectivity patterns of cerebellar neurons. We chose this method over spike-triggered-averaging maps (STM) ^24^, another method previously shown to reflect cortical connectivity patterns of subcortical neurons, primarily because of the difficulties it posed on computing average cortical images surrounding large numbers of spikes (e.g. high-firing Purkinje cells). To construct CorrMaps, spike trains were binned to match frame rate of wide-field images (40Hz) and convolved with the decay kernel of Ca^2+^ indicator GCaMP6s ^74^. Partial correlations were then calculated between convolved spike trains and each pixel time series with respect to the global cortical activity change (mean Δ*F/F* of all pixels over time), which were values shown in CorrMaps ^22^. The resulting CorrMaps were spatially smoothed using a 5x5 Gaussian filter (σ=3). To ensure the CorrMaps reflect stable, physiologically relevant functional connectivity relationship between the dorsal cortex and individual cerebellar neurons, we undertook the following three criteria:

1) **Criteria#1:** Number of spikes in the 10min recording session needed to exceed 200.
2) Significant correlation was determined by generating a null distribution of correlation values between each pixel time series and randomly permuted spike train (spike trains were divided into 1s blocks and shuffled 1000 times). All pixels with correlation values above 95th percentile or below 5th percentile of the null distribution were considered to be significantly positively or negatively correlated, respectively, with the spiking activity of a given cerebellar neuron. **Criteria#2:** The proportion of significantly correlated pixels needed to be greater than 0.3.
3) To ensure the CorrMaps were reproducible, interleaved 1s epochs of imaging and spiking data were concatenated and separately used to construct two separate CorrMaps; similarity (Pearson correlation) between these two interleaved epoch CorrMaps were compared to a null distribution generated using shuffled spike trains, and **Criteria#3:** CorrMaps with interleaved epoch similarity above 95% percentile of the null distribution were considered stable connectivity patterns.

Only cerebellar neurons with CorrMaps satisfying all three criteria in **both** anesthetized and awake conditions were included for further analysis.

#### Seed Pixel Correlation Map (SPM)

For each cortical imaging session, one SPM was calculated for each pixel (64x64 in total) by correlating the activity of a selected pixel with all other pixels. CorrMaps of individual cerebellar neurons were compared to the resulting library of SPMs of a given session to find the best matched SPM.

#### Parcellation of Dorsal Cortex

To facilitate comparing CorrMaps between animals and recording sessions, wide-field cortical images were parcellated and registered to Allen Mouse Brain Atlas (http://mouse.brain-map.org/) using a machine-learning based brain image segmentation tool MesoNet ^31^ (https://github.com/bf777/MesoNet). Briefly, landmarks on the skull (e.g. bregma, lambda) were automatically detected and boundaries of different brain regions were delimited using a pre-trained model specifically suited for mesoscale cortical images of C57BL/6 mice. Contours for each brain region detected by the algorithm were generated so that the pixel coordinates of each brain region on each CorrMap were obtained for subsequent analysis. This step was necessary for improved consistency in CorrMaps clustering.

#### Clustering of CorrMaps

To quantify the diversity of cerebellar CorrMaps, we performed clustering on all CorrMaps that passed the above quality control criteria. To facilitate comparison of CorrMaps in different states, CorrMaps obtained in anesthetized and awake states were clustered in one dataset. The high dimensional data (each CorrMap has 64x64=4096 pixels) was first dimensionally reduced and aligned by calculating activation index and deactivation index of each CorrMap ^32^. Activation/deactivation index of each parcellated cortical region were defined by the fraction of pixels in that region that were significantly positively correlated or negatively correlated (see Methods, quality control, criteria#2), respectively. These indexes were concatenated into a single vector (41 regions x2) that represent the overall pattern of each CorrMap which were used as input of the clustering algorithm.

We employed an unsupervised clustering algorithm, PhenoGraph^33^, to categorize CorrMaps. The graph was built in 2 steps: 1) it finds k nearest neighbors for N input vectors (activation/deactivation index) using Euclidean distance, resulting in N sets of k nearest neighbors, 2) a weighted graph is built where the edge weight between pairs of nodes depend on the number of neighbors they share. We then perform Louvain community detection ^75^ on this graph to partition the graph that maximizes modularity. This algorithm only requires one input parameter which is *k*, the number of nearest neighbors to be found for each input vector, and we selected the most stable number of clusters (Fig.2D) over the range of typically chosen values of *k* ^33^. We then performed a cluster identity refinement step, where individual CorrMap whose similarity (Peason correlation value) with group average CorrMap of the assigned cluster was lower than 0.4 was manually reassigned if its similarity with group average CorrMaps of any other clusters was higher than 0.4; otherwise, that CorrMap was excluded. A total of 519 out of 4606 CorrMaps (11.3%) were excluded by this criteria.

#### Cell-type Classification of Cerebellar Neurons

As a first pass, putative mossy fibers and Purkinje cells were identified using Phyllum (https://github.com/blinklab/phyllum) ^76^, a Phy2 plugin designed for celltype and layer identification in high-density recordings of the cerebellum. Manual curations were then performed where putative mossy fibers were confirmed based on their triphasic waveforms, short interspike interval and high firing irregularities, and putative Purkinje cells were confirmed based on high firing rates (>30Hz), regular firing patterns (harmonics in autocorrelogram) and presence of complex spikes ^39^.

#### Identification of Monosynaptically Coupled Cerebellar Neurons

To identify monosynaptic interactions between cerebellar neurons, we adopted a previously described method ^50^ studying cortical neurons. All possible pairs of cerebellar cortex neurons within a given recording were evaluated. To determine whether neuron A was synaptically coupled to neuron B, each spike in neuron B’s spike train was independently jittered randomly between -5 and 5 ms, and this process was repeated 1000 times to generate 1000 surrogate spike trains. Cross-correlogram between the original spike train of A and jittered surrogate spike trains of B were constructed. The threshold for excitatory/inhibitory interaction was calculated by adding/subtracting 2.6 times standard deviations (99% confidence interval) of max/min of each null distribution cross-correlogram from the average max/min of each cross-correlogram. We considered that there was a putative monosynaptic interaction between neuron A and B if the mean cross-correlation value of the real spike train A and B between +1ms and +4ms of crossed either the excitatory or inhibitory threshold (determined as above).

#### Identification of Cerebellar Neurons Responding to Sensory Stimulation

10ms bins of spike train data surrounding each stimulation event were extracted and averaged over 40 trials to construct peristimulus time histogram (PSTH) of each cerebellar neuron. PSTH was normalized by subtracting the mean value of the baseline period (500ms prior to stimulation onset). Response windows were between stimulus onset and 100ms or 300ms for facial stimulation and hindlimb stimulation respectively, and it was determined by the kinetics of firing rate changes to the stimulation (Fig.7B). Cerebellar neurons with means of their PSTH in the response window at least 2 standard deviations above/below the mean of the baseline period were considered stimulation responsive.

#### Statistics

Statistical tests were conducted in MATLAB. Data distributions were tested for normality using Kolmogorov-Smirnov tests prior to subsequent statistical analyses. All distributions were determined to be not normally distributed. Mann-Whitney U test and Wilcoxon signed-rank test were applied to determine difference between two unpaired and paired data distributions, respectively. Kolmogorov-Smirnov tests were applied to determine if two data groups were drawn from the same continuous distribution. Bootstrap test was applied to determine significance of pixel-to-spike-train correlation value, CorrMap stability (epoch correlation) and monosynaptic connections. All data was presented as mean±SEM unless specified otherwise.

## Acknowledgment

This work was supported by a Canadian Institutes of Health Research (CIHR) Foundation Grant FDN-143209 and project grant to T.H.M. THM was also supported by the Brain Canada Neurophotonics Platform, a Heart and Stroke Foundation of Canada grant in aid, National Science and Engineering Council of Canada (NSERC; GPIN-2022-03723), and a Leducq Foundation grant. This work was supported by resources made available through the Dynamic Brain Circuits cluster and the NeuroImaging and NeuroComputation Centre at the UBC Djavad Mowafaghian Centre for Brain Health (RRID SCR_019086) and made use of the DataBinge forum. We thank Pumin Wang, and Cindy Jiang for surgical assistance.

---

**Fig.S1.**
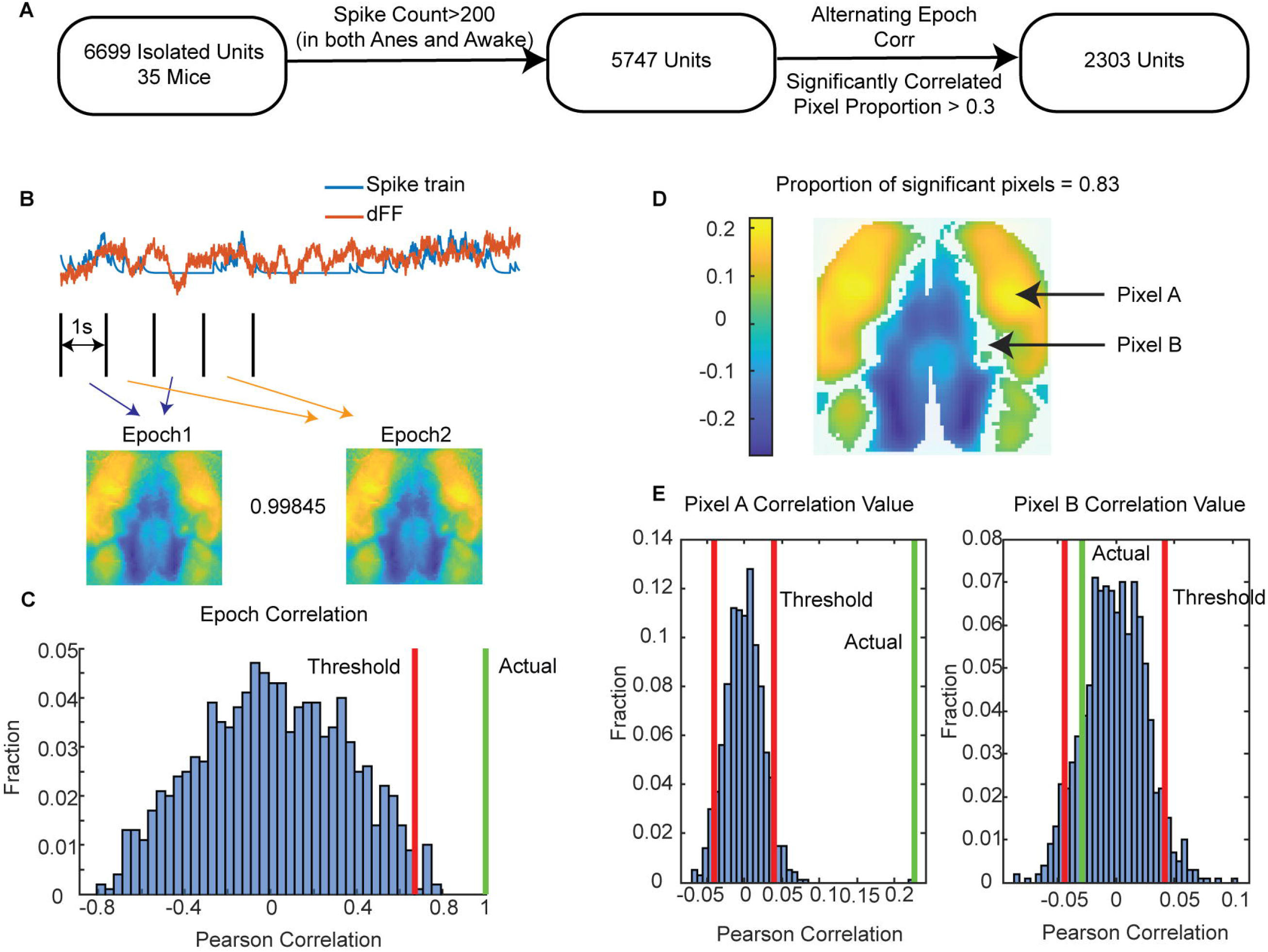
Not all cerebellar neurons show stable functional connectivity patterns with the dorsal cortex. A. Flow chart showing the 3 criteria used for identifying cerebellar neurons that have stable cortical representations. **B.** Convolved spike train and ΔF/F time series were split into 1s alternating epochs that were concatenated into epoch 1 and epoch 2 sequences. CorrMaps were generated separately for epoch 1 and epoch 2 sequences and the correlation between the two CorrMaps were termed epoch correlation. Epoch correlation for the example neuron shown is >0.99. **C.** Example null distribution histogram of epoch correlations generated using shuffled spike train and original imaging data (same neuron as in B). Only cerebellar neurons with epoch correlation greater than the 95% confidence threshold were included for analysis (Methods). **D.** Significance mask generated for CorrMap shown in B. **E.** Null distribution of pearson correlation value of pixel A and B calculated using shuffled spike train and original ΔF/F. Pixels with actual correlation value >95% or < 5% of the null distribution were considered significantly correlated (Methods).

**Fig.S2.**
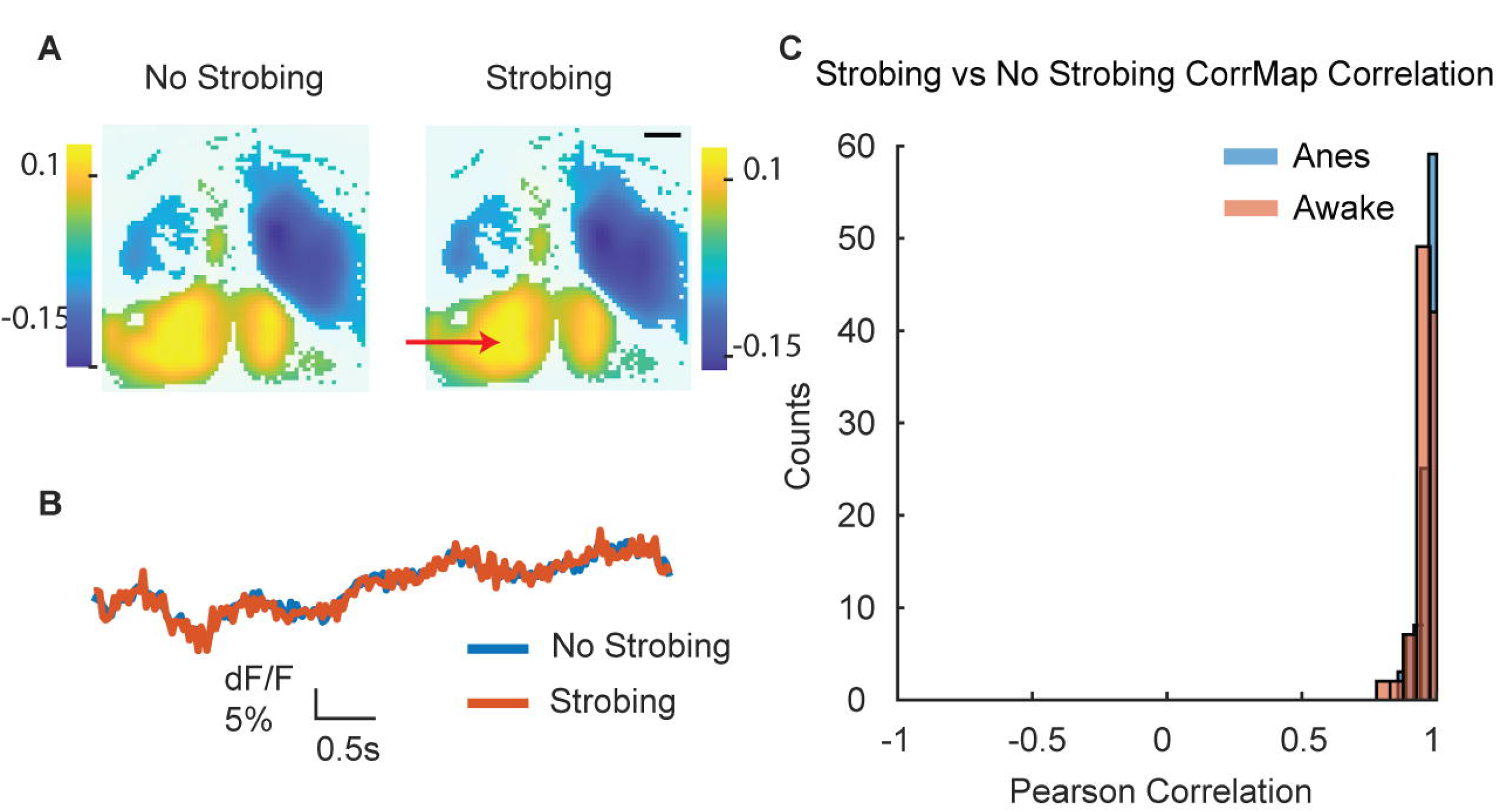
Hemodynamic correction does not affect the overall functional connectivity pattern. A. Example CorrMap calculated with or without hemodynamic correction using green reflected light strobing (Methods). Spatial scale bar-1mm. **B.** Example ΔF/F traces of brain activity with or without subtracting the hemodynamic green reflectance signal. **C.** Histogram of correlation values between CorrMaps of cerebellar neurons calculated with or without hemodynamic correction in anesthetized (blue) or awake (orange) state (anesthetized, 0.96±0.03; awake, 0.96±0.04; mean±std, n=102 neurons, 1 mouse).

**Fig.S3.**
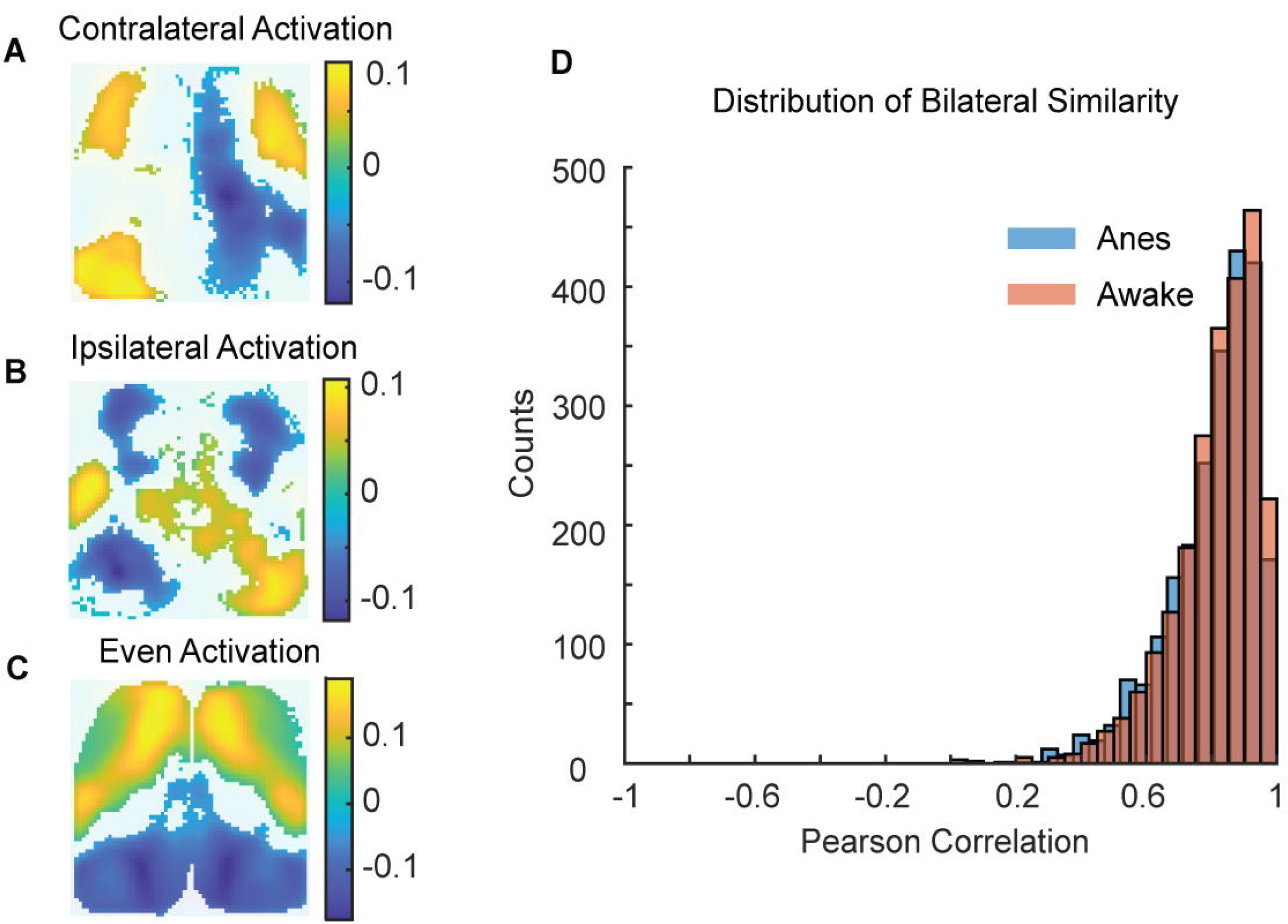
CorrMaps show high bilateral similarities A-B. Example CorrMaps of cerebellar neurons that show more activation in cortical regions contralateral (A) or ipsilateral (B) to the cerebellar recording site (cerebellar recordings were all made on the right side). **C.** Example CorrMap of a cerebellar neuron that showed even activation on both hemispheres. **D.** Histogram of similarity (Pearson correlation of concatenated activation and deactivation index vectors corresponding to the two hemispheres) of CorrMaps in anesthetized (blue) or awake (orange) state (anesthetized, 0.80±0.14; awake, 0.81±0.13; mean±std, n = 2303 neurons, 35 mice).

**Fig.S4.**
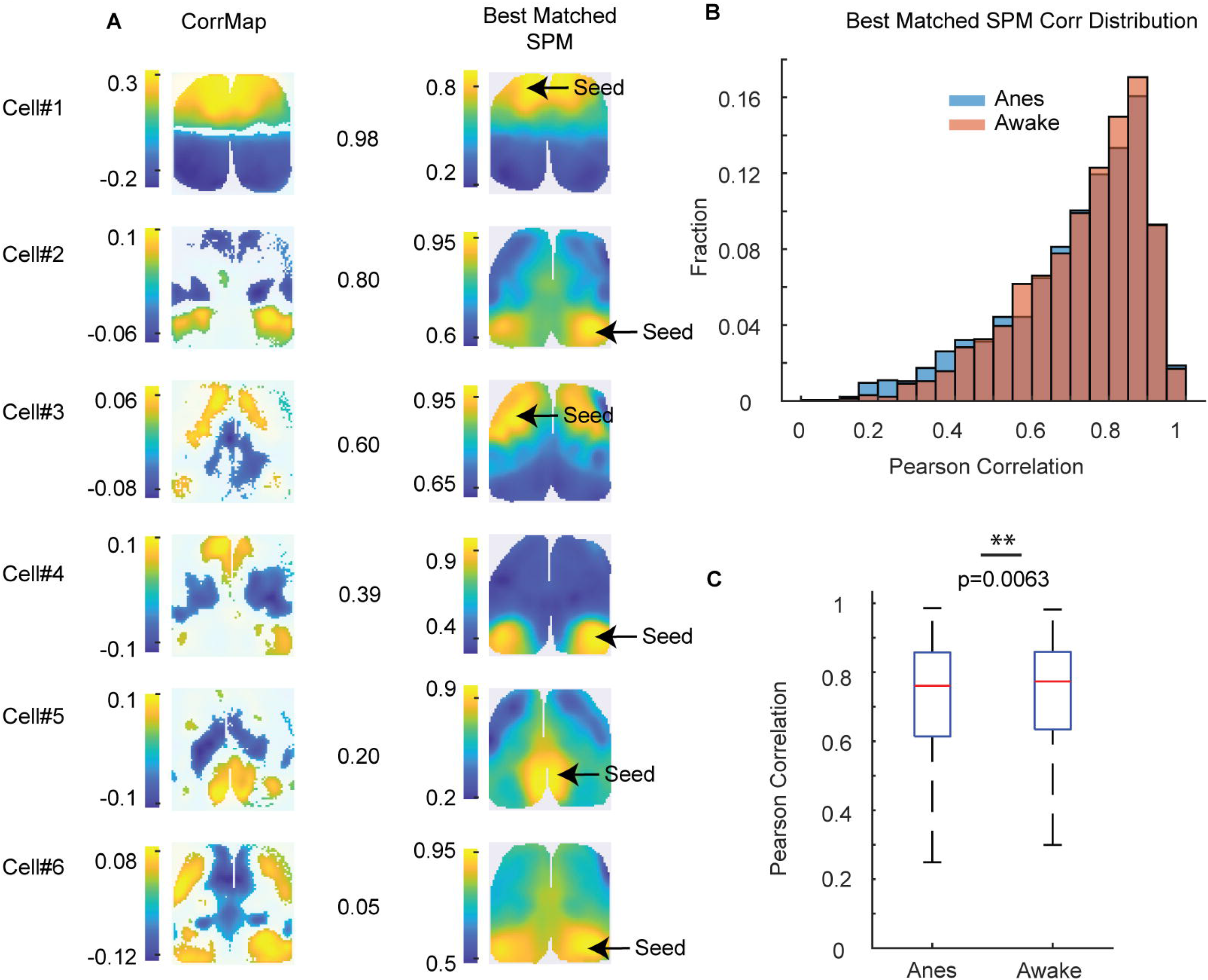
Functional connectivity patterns of most cerebellar neurons are consistent with intrinsic cortical activity motifs. A. Example CorrMaps and their similarities to their best-matching seed pixel correlation maps (SPM, see Methods). **B.** Histogram of the similarities (Pearson correlation value) between CorrMaps and their best-matching SPM in anesthetized (blue) or awake (orange) state. **C.** Average of the correlation values shown in B (anesthetized, 0.71±0.18; awake, 0.73±0.16, mean±std; Wilcoxon signed rank test, p=0.0063, N=2303 neurons, 35 mice).

**Fig.S5.**
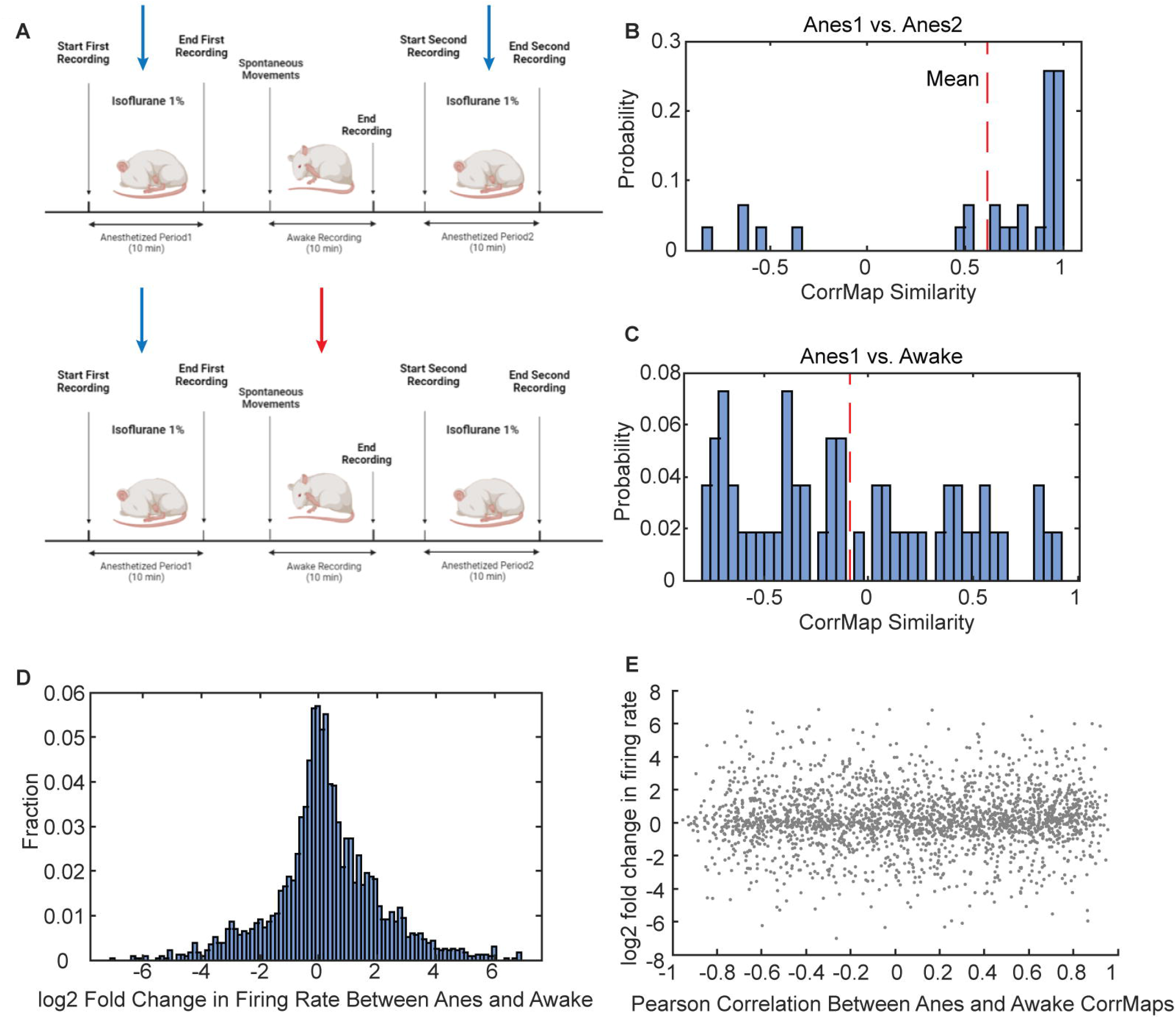
CorrMaps were stable between two consecutive anesthetized recordings. A. Schematic showing the sequence of state transitions in which the recordings were made. **B, C.** Histograms of the similarities (Pearson correlation value) between CorrMaps of the same cerebellar neurons recorded in the first and second anesthetized recording sessions (anesthetized session 1 vs anesthetized session 2 CorrMap similarity, 0.61±0.10, n = 4 mice, 31 neurons) or recorded in the anesthetized state first followed by the awake state (anesthetized session 1 vs awake session CorrMap similarity, -0.09±0.07, n = 4 mice, 55 neurons). **D.** Histogram of log2 fold change in firing rate between anesthetized and awake state for cerebellar neurons with stable CorrMaps (0.26±1.82, n=2303 neurons, 35 mice). **E.** Scatter plot of log2 fold change in firing rate against Pearson correlation of CorrMaps calculated in anesthetized and awake state. Each dot represents an individual cerebellar neuron (n=2303 neurons, 35 mice).

**Fig.S6.**
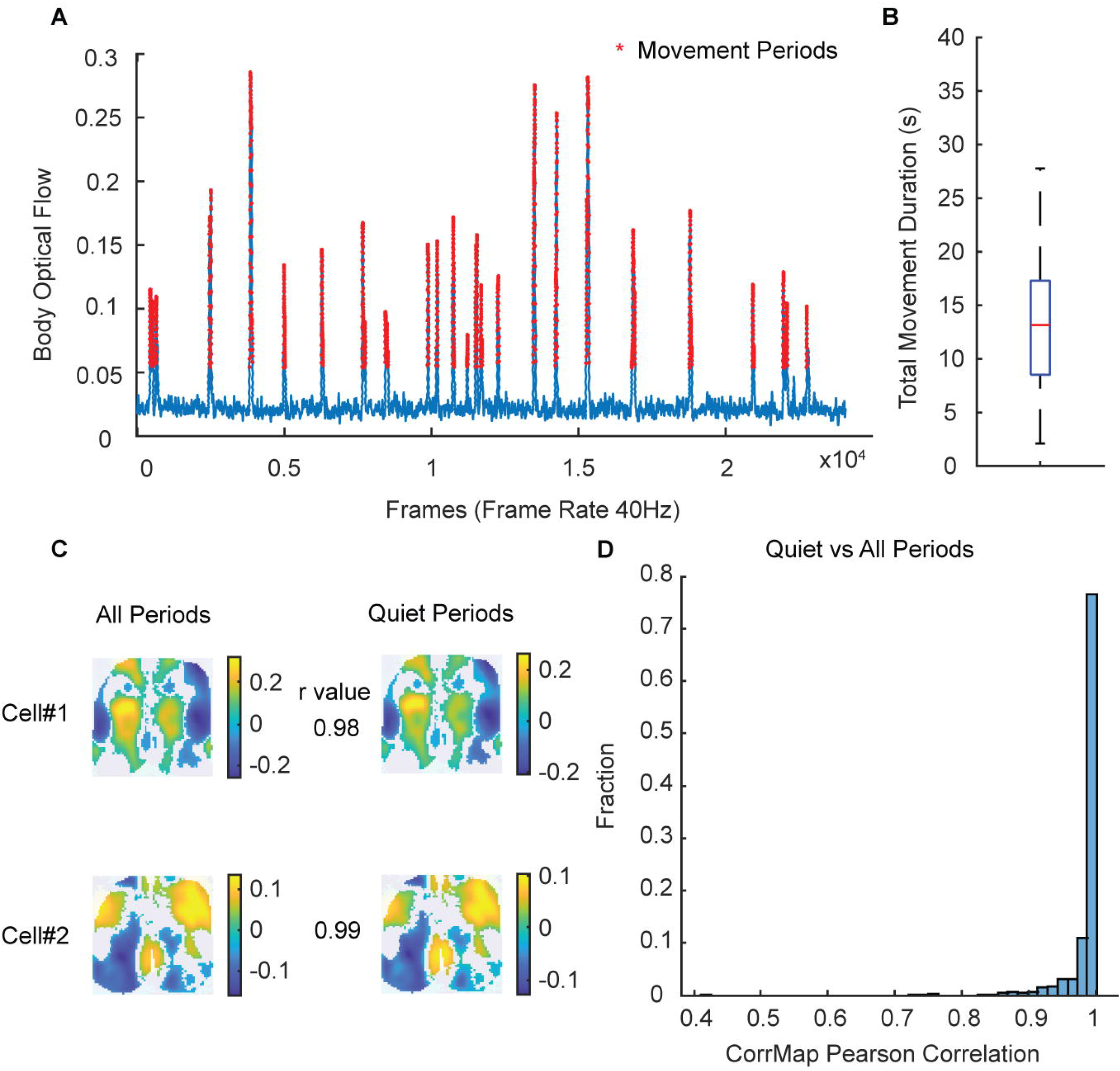
Spontaneous movements during awake recording have minimal impact on CorrMap patterns of cerebellar neurons. A. Spontaneous movement periods during awake state detected from behavioral videos (acquired at 40Hz) using frame-by-frame optical flow of manually defined animal body region of interest (ROI). **B.** Box plot of total movement duration for awake recordings (13.9±1.3s, 40 sessions from 10 mice, total recording duration was 600 s for each session). **C.** Example CorrMaps calculated using the entire recording or quiet periods (movement periods removed). **D.** Histogram of similarities (Pearson correlation) between CorrMaps of individual cerebellar neurons calculated using entire recording or quiet periods only (0.98±0.04, mean±std; 733 neurons from 10 mice).

## References

1. Ito, M. Mechanisms of motor learning in the cerebellum11Published on the World Wide Web on 24 November 2000. Brain Res. 886, 237–245 (2000).

2. Rochefort, C. et al. Cerebellum Shapes Hippocampal Spatial Code. Science 334, 385– 389 (2011).

3. Heffley, W. et al. Coordinated cerebellar climbing fiber activity signals learned sensorimotor predictions. Nat. Neurosci. 21, 1431–1441 (2018).

4. Kostadinov, D., Beau, M., Blanco-Pozo, M. & Häusser, M. Predictive and reactive reward signals conveyed by climbing fiber inputs to cerebellar Purkinje cells. Nat. Neurosci. 22, 950–962 (2019).

5. Wagner, M. J., Kim, T. H., Savall, J., Schnitzer, M. J. & Luo, L. Cerebellar granule cells encode the expectation of reward. Nature 544, 96–100 (2017).

6. Jackman, S. L. et al. Cerebellar Purkinje cell activity modulates aggressive behavior. eLife 9, e53229 (2020).

7. Carta, I., Chen, C. H., Schott, A. L., Dorizan, S. & Khodakhah, K. Cerebellar modulation of the reward circuitry and social behavior. Science 363, eaav0581 (2019).

8. Habas, C. et al. Distinct Cerebellar Contributions to Intrinsic Connectivity Networks. J. Neurosci. 29, 8586–8594 (2009).

9. Hoche, F., Guell, X., Sherman, J. C., Vangel, M. G. & Schmahmann, J. D. Cerebellar Contribution to Social Cognition. The Cerebellum 15, 732–743 (2016).

10. Stoodley, C. J. & Schmahmann, J. D. Functional topography in the human cerebellum: A meta-analysis of neuroimaging studies. NeuroImage 44, 489–501 (2009).

11. Schmahmann, J. D. & Sherman, J. C. The cerebellar cognitive affective syndrome. 19.

12. Wang, S. S.-H., Kloth, A. D. & Badura, A. The Cerebellum, Sensitive Periods, and Autism. Neuron 83, 518–532 (2014).

13. D’Angelo, E. & Casali, S. Seeking a unified framework for cerebellar function and dysfunction: from circuit operations to cognition. Front. Neural Circuits 6, 116 (2013).

14. Bloedel, J. R. Functional heterogeneity with structural homogeneity: How does the cerebellum operate? in Movement Control (eds. Cordo, P. & Harnad, S.) 64–76 (Cambridge University Press, 1994). doi:10.1017/CBO9780511529788.007.

15. Ramnani, N. The primate cortico-cerebellar system: anatomy and function. Nat. Rev. Neurosci. 7, 511–522 (2006).

16. Balsters, J. H. et al. Evolution of the cerebellar cortex: The selective expansion of prefrontal-projecting cerebellar lobules. NeuroImage 49, 2045–2052 (2010).

17. Swenson, R. S., Sievert, C. F., Terreberry, R. R., Neafsey, E. J. & Castro, A. J. Organization of cerebral cortico-olivary projections in the rat. Neurosci. Res. 7, 43–54 (1989).

18. Henschke, J. U. & Pakan, J. M. Disynaptic cerebrocerebellar pathways originating from multiple functionally distinct cortical areas. eLife 9, e59148 (2020).

19. Houck, B. D. & Person, A. L. Cerebellar Loops: A Review of the Nucleocortical Pathway. The Cerebellum 13, 378–385 (2014).

20. Kelly, R. M. & Strick, P. L. Cerebellar Loops with Motor Cortex and Prefrontal Cortex of a Nonhuman Primate. J. Neurosci. 23, 8432–8444 (2003).

21. Pisano, T. J. et al. Homologous organization of cerebellar pathways to sensory, motor, and associative forebrain. Cell Rep. 36, 109721 (2021).

22. Clancy, K. B., Orsolic, I. & Mrsic-Flogel, T. D. Locomotion-dependent remapping of distributed cortical networks. Nat. Neurosci. 22, 778–786 (2019).

23. Peters, A. J., Fabre, J. M. J., Steinmetz, N. A., Harris, K. D. & Carandini, M. Striatal activity topographically reflects cortical activity. Nature 591, 420–425 (2021).

24. Xiao, D. et al. Mapping cortical mesoscopic networks of single spiking cortical or sub-cortical neurons. eLife 6, e19976 (2017).

25. Jun, J. J. et al. Fully integrated silicon probes for high-density recording of neural activity. Nature 551, 232–236 (2017).

26. Madisen, L. et al. Transgenic mice for intersectional targeting of neural sensors and effectors with high specificity and performance. Neuron 85, 942–958 (2015).

27. Fox, M. D. et al. The human brain is intrinsically organized into dynamic, anticorrelated functional networks. Proc. Natl. Acad. Sci. 102, 9673–9678 (2005).

28. Bauer, A. Q. et al. Effective Connectivity Measured Using Optogenetically Evoked Hemodynamic Signals Exhibits Topography Distinct from Resting State Functional Connectivity in the Mouse. Cereb. Cortex 28, 370–386 (2018).

29. Mohajerani, M. H. et al. Spontaneous cortical activity alternates between motifs defined by regional axonal projections. Nat. Neurosci. 16, 1426–1435 (2013).

30. Vanni, M. P., Chan, A. W., Balbi, M., Silasi, G. & Murphy, T. H. Mesoscale Mapping of Mouse Cortex Reveals Frequency-Dependent Cycling between Distinct Macroscale Functional Modules. J. Neurosci. 37, 7513–7533 (2017).

31. Xiao, D., Forys, B. J., Vanni, M. P. & Murphy, T. H. MesoNet allows automated scaling and segmentation of mouse mesoscale cortical maps using machine learning. Nat. Commun. 12, 5992 (2021).

32. Barson, D. et al. Simultaneous mesoscopic and two-photon imaging of neuronal activity in cortical circuits. Nat. Methods 17, 107–113 (2020).

33. Levine, J. H. et al. Data-Driven Phenotypic Dissection of AML Reveals Progenitor-like Cells that Correlate with Prognosis. Cell 162, 184–197 (2015).

34. Buckner, R. L., Krienen, F. M., Castellanos, A., Diaz, J. C. & Yeo, B. T. T. The organization of the human cerebellum estimated by intrinsic functional connectivity. J. Neurophysiol. 106, 2322–2345 (2011).

35. King, M., Hernandez-Castillo, C. R., Poldrack, R. A., Ivry, R. B. & Diedrichsen, J. Functional boundaries in the human cerebellum revealed by a multi-domain task battery. Nat. Neurosci. 22, 1371–1378 (2019).

36. Diedrichsen, J., Hashambhoy, Y., Rane, T. & Shadmehr, R. Neural Correlates of Reach Errors. J. Neurosci. 25, 9919–9931 (2005).

37. Wagner, M. J. et al. A neural circuit state change underlying skilled movements. Cell 184, 3731–3747.e21 (2021).

38. Hull, C. & Regehr, W. G. The Cerebellar Cortex. Annu. Rev. Neurosci. 45, 151–175 (2022).

39. Dijck, G. V. et al. Probabilistic Identification of Cerebellar Cortical Neurones across Species. PLOS ONE 8, e57669 (2013).

40. Albus, J. S. A theory of cerebellar function. Math. Biosci. 10, 25–61 (1971).

41. Ito, M. Cerebellar Long-Term Depression: Characterization, Signal Transduction, and Functional Roles. Physiol. Rev. 81, 1143–1195 (2001).

42. Deverett, B., Koay, S. A., Oostland, M. & Wang, S. S.-H. Cerebellar involvement in an evidence-accumulation decision-making task. eLife 7, e36781 (2018).

43. Gruijl, J. R. D., Hoogland, T. M. & Zeeuw, C. I. D. Behavioral Correlates of Complex Spike Synchrony in Cerebellar Microzones. J. Neurosci. 34, 8937–8947 (2014).

44. Heffley, W. & Hull, C. Classical conditioning drives learned reward prediction signals in climbing fibers across the lateral cerebellum. eLife 8, e46764 (2019).

45. Gao, H., Solages, C. de & Lena, C. Tetrode recordings in the cerebellar cortex. J. Physiol.-Paris 106, 128–136 (2012).

46. Apps, R. & Garwicz, M. Anatomical and physiological foundations of cerebellar information processing. Nat. Rev. Neurosci. 6, 297–311 (2005).

47. Okun, M. et al. Diverse coupling of neurons to populations in sensory cortex. Nature 521, 511–515 (2015).

48. De Zeeuw. Bidirectional learning in upbound and downbound microzones of the cerebellum. Nat. Rev. Neurosci. 22, 92–110 (2021).

49. Valera, A. M. et al. Stereotyped spatial patterns of functional synaptic connectivity in the cerebellar cortex. eLife 5, e09862 (2016).

50. Fujisawa, S., Amarasingham, A., Harrison, M. T. & Buzsáki, G. Behavior-dependent short-term assembly dynamics in the medial prefrontal cortex. Nat. Neurosci. 11, 823– 833 (2008).

51. Vincent, J. L. et al. Intrinsic functional architecture in the anaesthetized monkey brain. Nature 447, 83–86 (2007).

52. Chadderton, P., Margrie, T. W. & Häusser, M. Integration of quanta in cerebellar granule cells during sensory processing. Nature 428, 856–860 (2004).

53. Eccles, J. C., Provini, L., Strata, P. & Táboříková, H. Analysis of electrical potentials evoked in the cerebellar anterior lobe by stimulation of hindlimb and forelimb nerves. Exp. Brain Res. 6, 171–194 (1968).

54. Morissette, J. & Bower, J. M. Contribution of somatosensory cortex to responses in the rat cerebellar granule cell layer following peripheral tactile stimulation. Exp. Brain Res. 109, 240–250 (1996).

55. Apps, R. & Watson, T. C. Cerebro-Cerebellar Connections. in Handbook of the Cerebellum and Cerebellar Disorders (eds. Manto, M., Schmahmann, J. D., Rossi, F., Gruol, D. L. & Koibuchi, N.) 1131–1153 (Springer Netherlands, 2013). doi:10.1007/978-94-007-1333-8_48.

56. Suzuki, L., Coulon, P., Sabel-Goedknegt, E. H. & Ruigrok, T. J. H. Organization of Cerebral Projections to Identified Cerebellar Zones in the Posterior Cerebellum of the Rat. J. Neurosci. 32, 10854–10869 (2012).

57. Liang, Z., Liu, X. & Zhang, N. Dynamic resting state functional connectivity in awake and anesthetized rodents. NeuroImage 104, 89–99 (2015).

58. Apps, R. et al. Cerebellar Modules and Their Role as Operational Cerebellar Processing Units. Cerebellum Lond. Engl. 17, 654–682 (2018).

59. Chen, X. et al. Transcriptomic mapping uncovers Purkinje neuron plasticity driving learning. Nature 605, 722–727 (2022).

60. Hull, C. Prediction signals in the cerebellum: Beyond supervised motor learning. eLife 9, e54073 (2020).

61. Ohmae, S. & Medina, J. F. Climbing fibers encode a temporal-difference prediction error during cerebellar learning in mice. Nat. Neurosci. 18, 1798–1803 (2015).

62. Fox, M. D. & Raichle, M. E. Spontaneous fluctuations in brain activity observed with functional magnetic resonance imaging. Nat. Rev. Neurosci. 8, 700–711 (2007).

63. Siegel, J. S. et al. Data Quality Influences Observed Links Between Functional Connectivity and Behavior. Cereb. Cortex N. Y. NY 27, 4492–4502 (2017).

64. Tsodyks, M., Kenet, T., Grinvald, A. & Arieli, A. Linking Spontaneous Activity of Single Cortical Neurons and the Underlying Functional Architecture. Science 286, 1943–1946 (1999).

65. McCormick, D. A., Nestvogel, D. B. & He, B. J. Neuromodulation of Brain State and Behavior. Annu. Rev. Neurosci. 43, 391–415 (2020).

66. Steinmetz, N. A. et al. Neuropixels 2.0: A miniaturized high-density probe for stable, long-term brain recordings. Science 372, eabf4588 (2021).

67. Dana, H. et al. Thy1-GCaMP6 Transgenic Mice for Neuronal Population Imaging In Vivo. PLOS ONE 9, e108697 (2014).

68. Siegle, J. H. et al. Open Ephys: an open-source, plugin-based platform for multichannel electrophysiology. J. Neural Eng. 14, 045003 (2017).

69. Pachitariu, M., Steinmetz, N. A., Kadir, S. N., Carandini, M. & Harris, K. D. Fast and accurate spike sorting of high-channel count probes with KiloSort. in Advances in Neural Information Processing Systems vol. 29 (Curran Associates, Inc., 2016).

70. Rossant, C. et al. Spike sorting for large, dense electrode arrays. Nat. Neurosci. 19, 634–641 (2016).

71. de Solages, C. et al. High-Frequency Organization and Synchrony of Activity in the Purkinje Cell Layer of the Cerebellum. Neuron 58, 775–788 (2008).

72. Ma, Y. et al. Wide-field optical mapping of neural activity and brain haemodynamics: considerations and novel approaches. Philos. Trans. R. Soc. B Biol. Sci. 371, 20150360 (2016).

73. Wang, Q. et al. The Allen Mouse Brain Common Coordinate Framework: A 3D Reference Atlas. Cell 181, 936–953.e20 (2020).

74. Chen, T.-W. et al. Ultrasensitive fluorescent proteins for imaging neuronal activity. Nature 499, 295–300 (2013).

75. Blondel, V. D., Guillaume, J.-L., Lambiotte, R. & Lefebvre, E. Fast unfolding of communities in large networks. J. Stat. Mech. Theory Exp. 2008, P10008 (2008).

76. Naveros, F., et al. Phyllum: a phy plugin for surveying high-density neural recordings in the cerebellum. in Program No. 299.05. 2022 Neuroscience Meeting Planner (2022).

